# Enhancers display constrained sequence flexibility and context-specific modulation of motif function

**DOI:** 10.1101/2022.08.31.506061

**Authors:** Franziska Reiter, Bernardo P. de Almeida, Alexander Stark

## Abstract

The information about when and where each gene is to be expressed is mainly encoded in the DNA sequence of enhancers, sequence elements that comprise binding sites (motifs) for different transcription factors (TFs). Most of the research on enhancer sequences has been focused on TF motif presence, while the enhancer syntax, i.e. the flexibility of important motif positions and how the sequence context modulates the activity of TF motifs, remain poorly understood. Here, we explore the rules of enhancer syntax by a two-pronged approach in *Drosophila melanogaster* S2 cells: we (1) replace important motifs by an exhaustive set of all possible 65,536 eight-nucleotide-long random sequences and (2) paste eight important TF motif types into 763 positions within 496 enhancers. These complementary strategies reveal that enhancers display constrained sequence flexibility and the context-specific modulation of motif function. Important motifs can be functionally replaced by hundreds of sequences constituting several distinct motif types, but only a fraction of all possible sequences and motif types restore enhancer activity. Moreover, TF motifs contribute with different intrinsic strengths that are strongly modulated by the enhancer sequence context (the flanking sequence, presence and diversity of other motif types, and distance between motifs), such that not all motif types can work in all positions. The context-specific modulation of motif function is also a hallmark of human enhancers and TF motifs, as we demonstrate experimentally. Overall, these two general principles of enhancer sequences are important to understand and predict enhancer function during development, evolution and in disease.

## Introduction

Transcriptional enhancers are DNA sequence elements that control gene expression by modulating the transcription of their target genes in specific cell types and conditions (Banerji et al. 1981; Levine 2010). These elements contain short sequence motifs bound by different transcription factors (TFs), and the combined regulatory cues of all bound TFs determine an enhancer’s activity (Spitz and Furlong 2012). Due to the critical role of enhancers in development, evolution and disease (Levine 2010; Rickels and Shilatifard 2018), understanding how enhancer sequences encode function is a major question in biology. Previous studies have highlighted the importance of sequence constraints within enhancers, such as the presence of TF motifs and features related to the motifs’ flanking sequences, affinities, and arrangements (their number, order, orientation and spacing), termed here ‘motif syntax’ (King et al. 2020; Jindal and Farley 2021; Ludwig et al. 2000; Kulkarni and Arnosti 2003; Zinzen et al. 2006; Panne 2008; Swanson et al. 2010; Liu and Posakony 2012; Erceg et al. 2014; Crocker et al. 2015; Farley et al. 2016, 2015; Fiore and Cohen 2016; Smith et al. 2013; Sharon et al. 2012; Thanos and Maniatis 1995; Hanes et al. 1994; Arnosti et al. 1996; Avsec et al. 2021; de Almeida et al. 2022). However, while mutations in enhancer sequences can change enhancer function and lead to morphological evolution and disease (Visel et al. 2009; Levine 2010; Gompel et al. 2005; Rickels and Shilatifard 2018), enhancers usually display only modest or no sequence conservation across species (Villar et al. 2015; May et al. 2012; Blow et al. 2010; Schmidt et al. 2010; Arnold et al. 2014; Fuqua et al. 2020; Ludwig et al. 1998) and even random DNA sequences can show enhancer activity (Galupa et al. 2022; de Boer et al. 2019). Therefore, the importance of sequence constraints and motif syntax within enhancers remain outstanding questions in gene regulation.

Two main models have been proposed to explain how enhancer sequence relates to function. The *enhanceosome* model assumes very strict syntax rules with invariant motif arrangements required for cooperative TF binding (Panne 2008). In contrast, the *billboard* model proposes that TFs bind independently without constraints on how motifs are arranged within the enhancer (Arnosti and Kulkarni 2005; Kulkarni and Arnosti 2003). Yet very few enhancers fit these models, having either invariant syntax or no constraints at all, and most enhancers fall in between these two extremes, with a flexible syntax yet high degree of dependency between enhancer features (Kulkarni and Arnosti 2003; Vockley et al. 2017; Jindal and Farley 2021). This complexity in enhancer sequence has prevented the generalization of sequence-rules derived from individual enhancers into unifying principles of the regulatory code, thus limiting our understanding of the sequence constraints related to motif syntax and TF activity in enhancers.

Although enhancer sequences evolve rapidly, their function can be conserved despite significant sequence changes (Rastegar et al. 2008; Ludwig et al. 1998, 2000; Swanson et al. 2011; Taher et al. 2011; Fisher et al. 2006; He et al. 2011; Wong et al. 2020; Weirauch and Hughes 2010; Farley et al. 2015; Blow et al. 2010; Schmidt et al. 2010; May et al. 2012; Villar et al. 2015; Arnold et al. 2014; Vaishnav et al. 2022). This suggests that there is considerable flexibility within enhancer sequences, and that the maintenance of function-defining features rather than overall sequence similarity is important for enhancer activity. This is illustrated most clearly by the maintenance of TF motifs at invariant positions or at different relative positions within orthologous enhancer sequences (Ludwig et al. 1998, 2000; Arnold et al. 2014; Wong et al. 2020; Rastegar et al. 2008). However, how flexible or constrained motif positions within enhancers are at both, the DNA sequence and the TF motif level, i.e. how many different sequence variants or motif types might functionally replace the wildtype sequence at important motif positions, has remained unknown. Similarly, even though TF motifs have been observed to move between different enhancer positions over the course of evolution (presumably a consequence of motif decay and de novo formation), and despite position-independence being a key assumption of the billboard model, the influence of the position and sequence context on a motif’s contribution to enhancer function is not understood. These knowledge gaps restrict our understanding of the functional and evolutionary flexibility of enhancer sequences and how many sequence variants, as they might arise by DNA mutagenesis, might lead to similar or different enhancer activities.

Here, we investigated how many defined DNA sequences might functionally replace the wildtype sequence in various motif and control positions by exhaustively testing all possible 8-nucleotide-long random sequence variants at these positions in two enhancers in *Drosophila melanogaster* S2 cells. At each position, hundreds of sequence variants corresponding to several different motif types could functionally replace the wildtype sequence (i.e. constitute *solutions*), suggesting that enhancer sequences display flexibility within and across motif types. However, at each position, these solutions constituted only a tiny fraction of the approximately 65,000 possible sequences, indicating that enhancer sequence flexibility is constrained. In addition, the solutions differed between positions and most TF motifs had highly context-dependent activities. Indeed, across hundreds of enhancer positions eight prominent motif types contributed to enhancer activity with different intrinsic strengths that were further modulated by the enhancer sequence context, namely the flanking sequence, the presence and diversity of other motif types, and the distance between motifs. The modulation of TF motif activity by the sequence context is a general enhancer feature, as we also demonstrate in human cells.

## Results

### STARR-seq comprehensively assesses the activity of enhancer variants revealing constrained enhancer sequence flexibility

To systematically test what sequences function in a certain enhancer position, we used an approach inspired by studies that tested the activity of fully randomized regulatory sequences (Farley et al. 2015; Galupa et al. 2022; Vaishnav et al. 2022; de Boer et al. 2019) or the local fitness landscape of the green-fluorescent protein (GFP; (Sarkisyan et al. 2016; Somermeyer et al. 2022)). We generated a comprehensive library of sequence variants by replacing a specific 8nt stretch of an enhancer with randomized nucleotides (N_8_) and assessed the enhancer activity of each variant by UMI-STARR-seq in *Drosophila* S2 cells (Fig 1A; see Methods; (Arnold et al. 2013; Neumayr et al. 2019)). We tested the power of this approach in the position of a GATA TF motif within the *ced-6* enhancer (*ced-6* position 241nt, or *pos241*) that is required for its activity. We recovered all possible 8nt variants (65,536) in the input library and obtained reliable enhancer activity measurements for each variant (Fig S1). This showed that the vast majority of all variants drive low activity levels, while only 374 (<1%) achieve similar activity to wildtype (+/-10%) and 600 (1%) drive even higher activity, i.e. constitute valid *solutions* at this motif position (Fig 1B).

**Figure 1.**
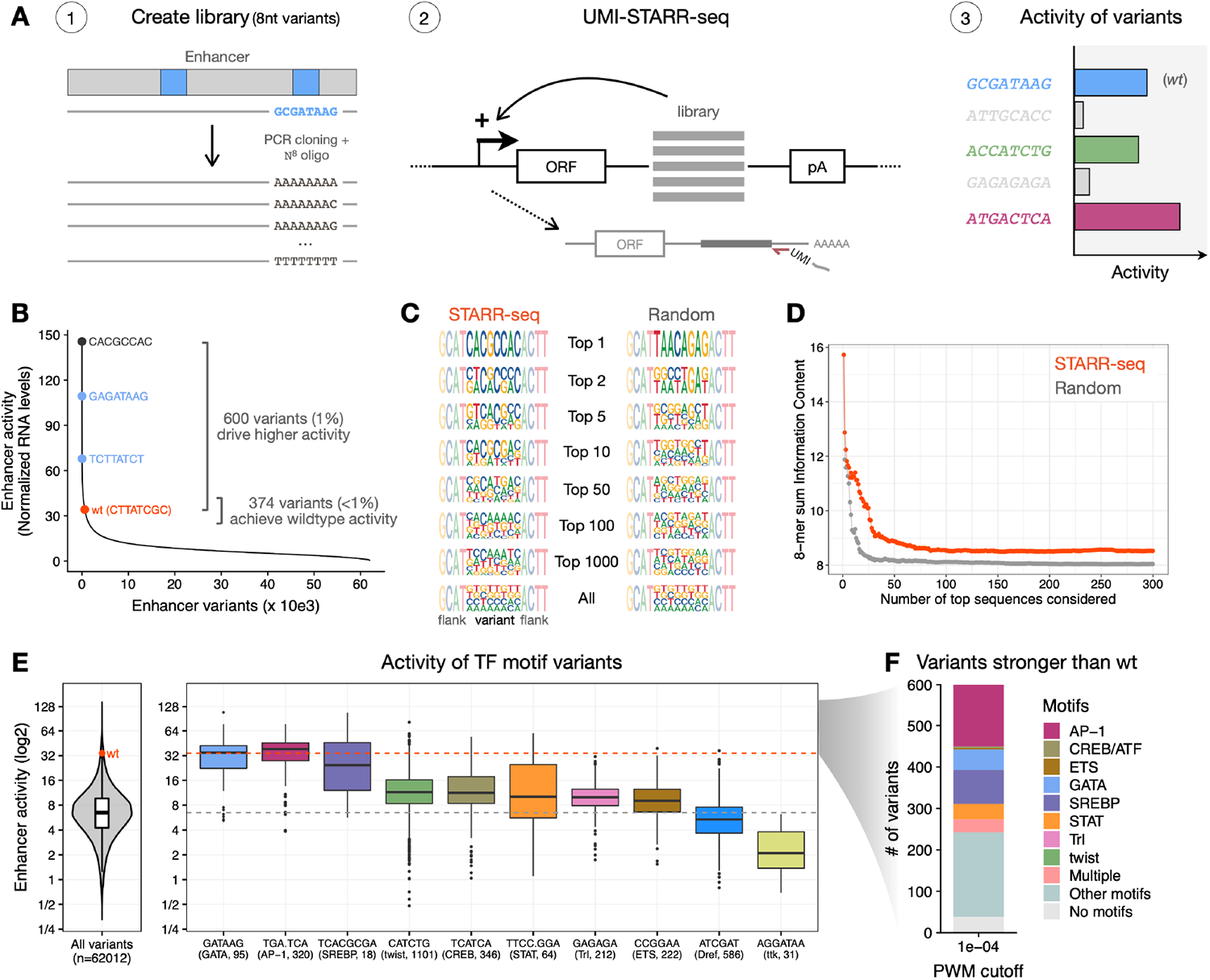
STARR-seq comprehensively assesses the activity of random variants in a specific enhancer position. **A)** Schematics of STARR-seq for the analysis of random variants in an enhancer position: (1) a comprehensive library of sequence variants was generated by replacing the 8nt stretch overlapping a GATA TF motif in the strong *ced-6* enhancer with all possible 65,536 randomized nucleotides; (2) the enhancer activity of each variant was measured by STARR-seq in *Drosophila* S2 cells; (3) expected outcomes include the wildtype sequence (wt, blue), inactive variants (grey), and variants that recover the wildtype activity (green) or are even stronger (purple). **B)** Most sequence variants exhibit low activity levels. The distribution of enhancer activity for each of the 62,012 enhancer variants with confident activity is shown. The wildtype (wt, red) sequence, the strongest GATA variant in each orientation (blue) and the strongest sequence variant are highlighted, together with the number of variants that achieve similar activity to wildtype (+/-10%) or drive even higher activity. **C)** Strong sequence variants are highly diverse. Logos with nucleotide frequency of the most-active variants in STARR-seq (1, 2, 5, 10, 50, 100, 1,000 and all; red). These were compared with the same logos after randomly sorting the variants (grey). **D)** Sum of information content within the most-active 8-mers in STARR-seq (red) compared with the same after randomly sorting the variants (grey), considering different number of top sequences. **E)** Distribution of enhancer activity for all 62,012 enhancer variants (left) or variants creating each TF motif (right). The activity of the wildtype sequence (wt, red dot and dashed line) or median of all variants (grey dashed line) are shown. The string of each TF motif used for the motif matching and the number of variants matching to each motif are described in the x-axis in the format “motif string (TF motif name, number of variants)”. **F)** Number of variants among the 600 stronger than wildtype that match to motifs enriched in S2 developmental enhancers (PWM p-value cutoff 1e^-04^).

Although only a few hundred sequences functioned at this position, these were highly diverse (Fig 1C,D) and included not only different variants of the GATA motif (Fig 1B – in blue, and 1E,F) but also other TF motifs, such as SREBP and AP-1 (Fig 1E,F, S2A,B, S3A). Indeed, most of the 600 variants stronger than wildtype (94%) created TF motifs overrepresented in S2 developmental enhancers (PWM p-value 1e^-04^; Fig 1F, S3B), showing that there is flexibility in the DNA sequences but also in the motif types they encode. However, different TF motifs rescued enhancer activity to different levels (Fig 1E, S3A). While AP-1 and SREBP achieved similar activity to the wildtype GATA motif, twist and ETS had lower activity at this enhancer position, despite being generally associated with strong enhancer activity in S2 cells (de Almeida et al. 2022). Therefore, the observed sequence flexibility is constrained to some TF motifs. In addition, even within each TF motif not all specific sequence variants functioned similarly, as apparent in the large differences between their activities (Fig 1E).

We also observed TF motif types that had neutral or repressive functions at the tested 8nt position: The Dref motif, previously shown to only be important for housekeeping enhancers (Zabidi et al. 2015; de Almeida et al. 2022), had no activity in this *ced-6* developmental enhancer, while the Ttk motif created the most inactive 8nt variants consistent with Ttk’s function as a repressor (Fig 1E, S2C; (Xiong and Montell 1993)). These results show that this approach can comprehensively assess the activity of all random variants in a specific region of the enhancer and identify activating, neutral and repressive sequences. Moreover, our findings indicate that developmental enhancers exhibit *constrained flexibility*, in that many variants, but still a strongly restricted number, can function at a given enhancer position. This constrained sequence flexibility applies not only to individual DNA sequences but also TF motif types in that several different motif types work, but not many or all.

### Activity of random variants in seven specific positions of two different enhancers

To evaluate if the same principles and the same specific solutions apply at different enhancer positions, we selected three additional positions of the *ced-6* enhancer and three positions of a strong enhancer in the *ZnT63C* locus (Fig 2A). To probe enhancer sequence flexibility at important motif positions and non-important control positions, we used the deep learning model DeepSTARR (Fig 2A; (de Almeida et al. 2022)) and previous experimental enhancer mutations (Fig S4F) to choose positions that should (*ced-6* pos110, pos241; *ZnT63C* pos142, pos180, pos210) or should not (*ced-6* pos182, pos230) be important for enhancer activity. We generated exhaustive libraries of all 8nt sequence variants for each position and performed UMI-STARR-seq on the combined libraries of each enhancer (S4A-E; see Methods). As observed for the GATA position in Fig 1 (pos241), only a restricted set of variants achieved wildtype activity at a second important GATA motif position in the same enhancer (pos110) or at the important motif positions in the *ZnT63C* enhancer (Fig 2B), confirming that important positions in enhancers show constrained flexibility. This contrasted with the non-important positions (pos182 and pos230 of the *ced-6* enhancer) where most sequence variants were active at wildtype levels or above (Fig 2B). Thus, the importance of an enhancer position reflects its constraint, with non-important positions not being constrained (while they can still be modulated positively or negatively).

**Figure 2.**
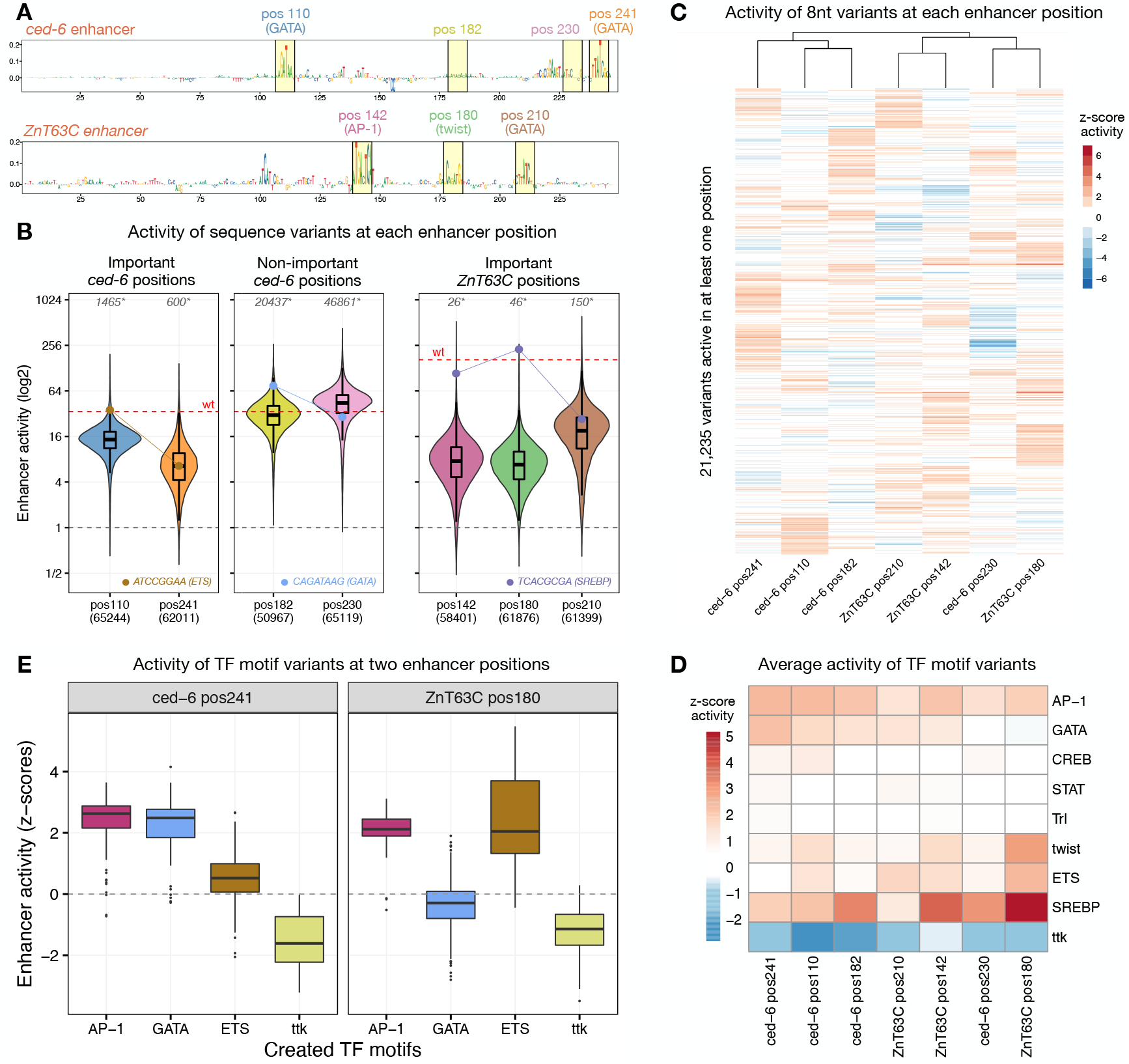
Sequence constraints at different enhancer positions. **A)** DeepSTARR-predicted nucleotide contribution scores for the *ced-6* (top) and *ZnT63C* (bottom) selected enhancer sequences. Selected 8nt motif positions and non-important control positions are highlighted in yellow with the respective numerical position, TF motif identity and different colors. **B)** Distribution of enhancer activity for all enhancer variants detected in each enhancer position. The activity of the wildtype sequence of each enhancer (wt, red dashed line) or of inactive sequences (grey dashed line) are highlighted, together with the activity of example sequence variants that create different TF motifs (ETS, GATA and SREBP; dots and connected lines). Number of variants tested in each position are shown in the x-axis, while the number of variants with higher activity than wildtype is shown on the top (grey, *). **C)** Heatmap of z-scores of log2 enhancer activity of 21,235 variants across all seven enhancer positions. Only variants assessed in all positions and active (z-score > 1) in at least one are shown. Variants were clustered using hierarchical clustering and their activity is colored in shades of red (activating) and blue (repressing). **D)** Heatmap of average z-scores of log2 enhancer activity of variants creating each TF motif type (y-axis) across all enhancer positions (x-axis; sorted as in (C)). Motif activity is colored in shades of red (activating) and blue (repressing). **E)** Distribution of z-scores of log2 enhancer activity for variants creating each of four TF motifs (AP-1, GATA, ETS, ttk) in two selected enhancer positions (*ced-6* pos241 and *ZnT63C* pos180).

The most active sequences at each enhancer position were highly diverse and exhibited distinct nucleotide preferences (Fig S5, S6). For example, two positions located either in the *ced-6* (pos110) or the *ZnT63C* (pos210) enhancer showed distinct preferences among the strongest 100 variants, which preferentially match to an SREBP (GTCAC[flanked by GTC]) or an ETS motif (CCGGA[A]), respectively (Fig S5B). These results show that different enhancer positions require different motif types and thus are under different constraints.

### Different TF motif types are active at different enhancer positions

Comparing the activity of the 8nt sequence variants between the enhancer positions (scaled to the average activity of variants to be comparable across positions; see Methods) revealed that they indeed functioned differently at different positions (Pearson correlation coefficients (PCCs) below 0.4 between positions; Fig 2C, S7A-C). Further consolidating the 8nt into 6nt variants to reduce the impact of the surrounding sequence of each position (averaged activity across the flanking nucleotides) still showed similar results (S7A,D). The top variants and solutions of each position differed substantially, with each position revealing specific sequences with particularly high activity, matching to known TF motifs (Fig 2C). For example, an ETS motif variant was amongst the strongest sequences at *ced-6* pos110 but not at pos241, a GATA variant was very active at *ced-6* pos182 but inactive at pos230, and a SREBP variant was active in all positions of the *ZnT63C* enhancer except at pos210 (Fig 2B).

We next compared the activity of motifs between the seven positions of the two different enhancers, by consolidating the activity of all 8nt variants (+/-4nt flanks) creating each motif (Fig 2D,E, S8; see Methods). Importantly, for each position the wildtype sequence as well as different variants of that motif were among the top variants. While the repressor ttk motif repressed in all positions and showed little specificity (similar to other known and novel repressor motifs; Fig S9), the activator motifs showed distinct profiles, such as motifs that are globally active in all positions (AP-1), motifs with low activity in all tested positions (STAT, CREB and Trl) and motifs with highly context-dependent activities (GATA, twist, ETS and SREBP) (Fig 2D,E). For example, GATA was active at the *ced-6* pos110 but not at the *ZnT63C* pos180 position, whereas ETS motifs showed the opposite profile with the strongest activity at *ZnT63C* pos180 (Fig 2E). Interestingly, for GATA motifs we observed strong activity in all positions except on *ced-6* pos230 and *ZnT63C* pos180, which are positioned close to another GATA motif (Fig 2A). This observation is in line with the previously observed negative interaction of GATA/GATA motif pairs at short distances (de Almeida et al. 2022) and suggests that the observed different activities of TF motifs at different enhancer positions depend on their interaction with other TFs and the sequence context.

In summary, testing thousands of random variants in different enhancer positions revealed that enhancer sequences display constrained flexibility, in that only a specific but still diverse set of sequences and TF motifs can function at a given position. However, importantly, these constraints and solutions differed between enhancer positions, with different TF motifs active at different positions, suggesting that their activity is modulated by the sequence context.

### Systematic motif pasting shows that motifs work differently at different enhancer positions

To systematically test if and how the enhancer sequence context modulates the function of TF motifs, we selected eight TF motifs that showed distinct position-dependent preferences (GATA, Trl, SREBP, AP-1, Atf2, twist, Stat92E and ETS) and pasted their optimal sequences into 763 positions in a total of 496 enhancers (Fig 3A; see Methods). These positions were selected to be TF motifs important for the activity of the respective enhancers, as assessed by motif mutagenesis, allowing the reliable measurement of the increase in enhancer activity after pasting each TF motif (here quantified as the log2 fold-change activity over the motif-mutated enhancer). UMI-STARR-seq experiments with these designed libraries produced highly reproducible and quantitative enhancer activity measurements (replicates PCC between 0.95 and 0.98; Fig S10A,B). Disrupting the selected enhancer positions by shuffling the wildtype sequences substantially reduced the activity of the respective wildtype enhancers by an average of more than 6-fold, and pasting the different TF motifs in these same positions rescued enhancer activity to different levels (Fig S10C). Since for each TF motif we pasted the same optimal sequence into all positions, the differences in activity can only be explained by their respective sequence context; the differences between TF motifs are also directly comparable, since we pasted them in the same set of positions.

**Figure 3.**
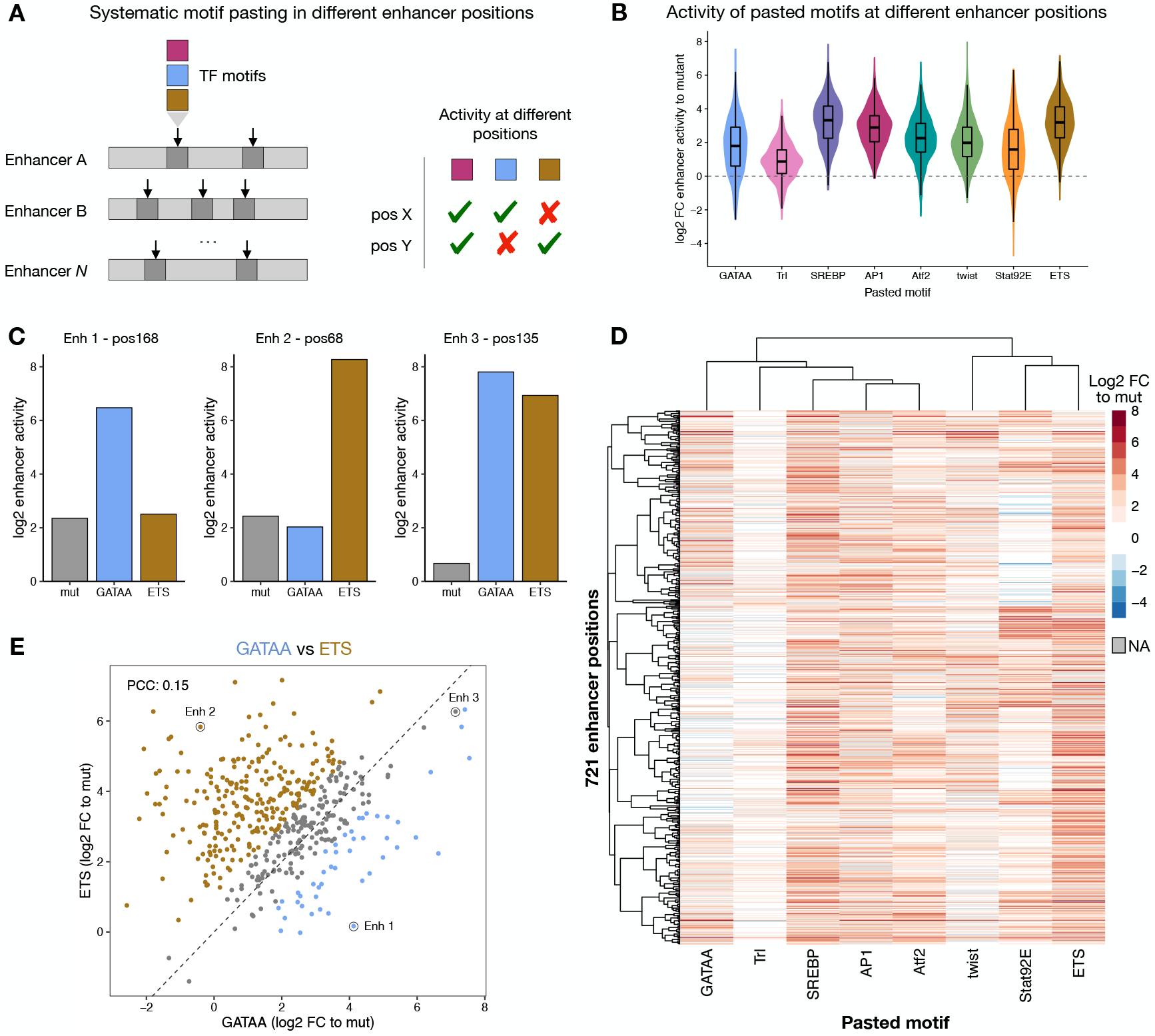
TF motifs work differently at different enhancer positions. **A)** Schematics of systematic motif pasting in different enhancer positions. Eight TF motifs that showed distinct position-dependent preferences were selected and their optimal sequence was pasted in 763 positions distributed among 496 enhancers, representing different contexts. The enhancer activity of each variant was measured in STARR-seq in *Drosophila* S2 cells to quantify the activity of motifs at the different positions. **B)** Distribution of enhancer activity changes (log2 FC to mutated sequence) across all enhancer positions for each pasted TF motif. **C)** Bar plots with activity (log2) of variants of three different enhancers with a mutated sequence (grey), a GATA (blue) or a ETS (brown) motif pasted at the same position. **D)** Heatmap of enhancer activity changes (log2 FC to mutated sequence) after pasting each of the eight selected TF motifs in 721 enhancer positions (positions with data for at least six motifs). TF motifs and positions were clustered using hierarchical clustering and the activity is colored in shades of red (activating) and blue (repressing); missing values are colored in grey. **E)** GATA and ETS motifs work differently at different enhancer positions. Comparison between enhancer activity changes (log2 FC to mutated sequence) after pasting GATA (x-axis) or ETS (y-axis) across all enhancer positions. Positions with stronger activity of GATA or ETS (>= 2-fold in respect to the other motif) are colored in blue and brown, respectively. Enhancer positions shown in (C) are highlighted. PCC: Pearson correlation coefficient.

Across all positions TF motifs had different median activities, which we interpret as different *intrinsic strengths*, with SREBP, ETS and AP-1 being the strongest and Trl the weakest motifs (Fig 3B, S10C). However, enhancer positions had large effects on the motif activities that differed more than 100-fold for the same motif (Fig 3B). For example, pasting a GATA motif activated enhancer activity more than 20-fold for 33 positions but not at all for 72 different positions. This position-dependency was particularly strong for Trl, Stat92E and GATA motifs, and weaker for AP-1, SREBP and ETS (Fig S10D), which all had higher intrinsic strengths. Additionally, each TF motif showed differential activity across enhancer positions and activated in a unique set of positions. For example (Fig 3C), GATA motifs activated enhancer1-position168 but not enh2-pos68, while ETS showed the opposite effect, and both motifs activated enh3-pos135. The different TF motifs showed different activity profiles across all positions, as revealed by global comparisons and hierarchical clustering (Fig 3D, S11). For example, GATA showed differential activity from ETS (PCC=0.15; Fig 3E) or twist (PCC=0.33; Fig S11B), while others such as AP-1 and Atf2 showed more similar positional preferences (PCC=0.68; Fig S11). These results highlight the complexity of enhancers syntax and the difficulty of predicting and interpreting individual sequence manipulations.

The distinct preferences observed between pasted motifs were largely independent of the identity of the replaced wildtype motif across all positions, as revealed by the weak interaction scores between the wildtype and the pasted motif identity in a multivariate linear regression analysis of all motif-pasting experiments (< 1% explained variance, Fig S12). In contrast, the pasted motif identity (irrespective of the identity of the replaced motif) explains the most (23%) while 65% of variance remains unexplained and is likely due to surrounding enhancer sequence features affecting the motifs’ activities. Thus, systematic pasting of TF motifs across hundreds of enhancer contexts shows that motifs have different intrinsic strengths but work differently at different enhancers and positions, suggesting that the enhancer sequence context constrains the activity of TF motifs.

### TF motifs have different intrinsic strengths that are modulated by the enhancer sequence context

The observed differential activities of motifs in different enhancer positions (Fig 3D) suggests that the enhancer sequence context modulates the function of TF motifs. We found no significant differences when comparing the motif activity between pairs of positions in the same enhancer or in different enhancers, suggesting that the local context immediately surrounding the motif is as important as enhancer identity (Fig S13).

More globally, the sequence context for a motif can be related to its position within the enhancer, the motif flanking sequence and the presence and distance to other motifs. To characterize the importance of these features, we tested if they contribute to the performance of predicting enhancer activity following the pasting of a motif at different enhancer positions. We first built a baseline random forest model that only includes the importance of the wildtype motif and the identity of the wildtype and pasted motifs as features, thereby not taking any sequence context features into account. This model obtained a PCC of 0.59 in the whole dataset using 10-fold cross-validation and showed that the pasted motif and the wildtype motif importance are strong determinants for enhancer activity (Fig S14A). Training a second random forest model that also includes context features such as the motif position relative to the enhancer center, the motif flanking sequence, and the presence and distance to other TF motifs, improved this performance to a PCC of 0.69 (Fig S14B). This shows that the enhancer sequence context, particularly the closest flanking nucleotides as well as the presence of other motifs at specific distances (e.g. GATA or ETS) have an impact on the activity of TF motifs (Fig S14B).

To better characterize the importance of these sequence rules for each TF motif separately, we generated interpretable linear models based on these rules to predict the motif activities across all positions (Fig 4A). These models were able to predict the motif pasting results, with PCCs to experimentally assessed log2 fold-changes between 0.39 (ETS) and 0.64 (Stat92E) (Fig 4A, S15). The motif flanks and the presence of additional motifs explained on average 16.7% and 6.7% of the motif activities variance, respectively, while the motif position within the enhancer had lower importance (0.4%).

**Figure 4.**
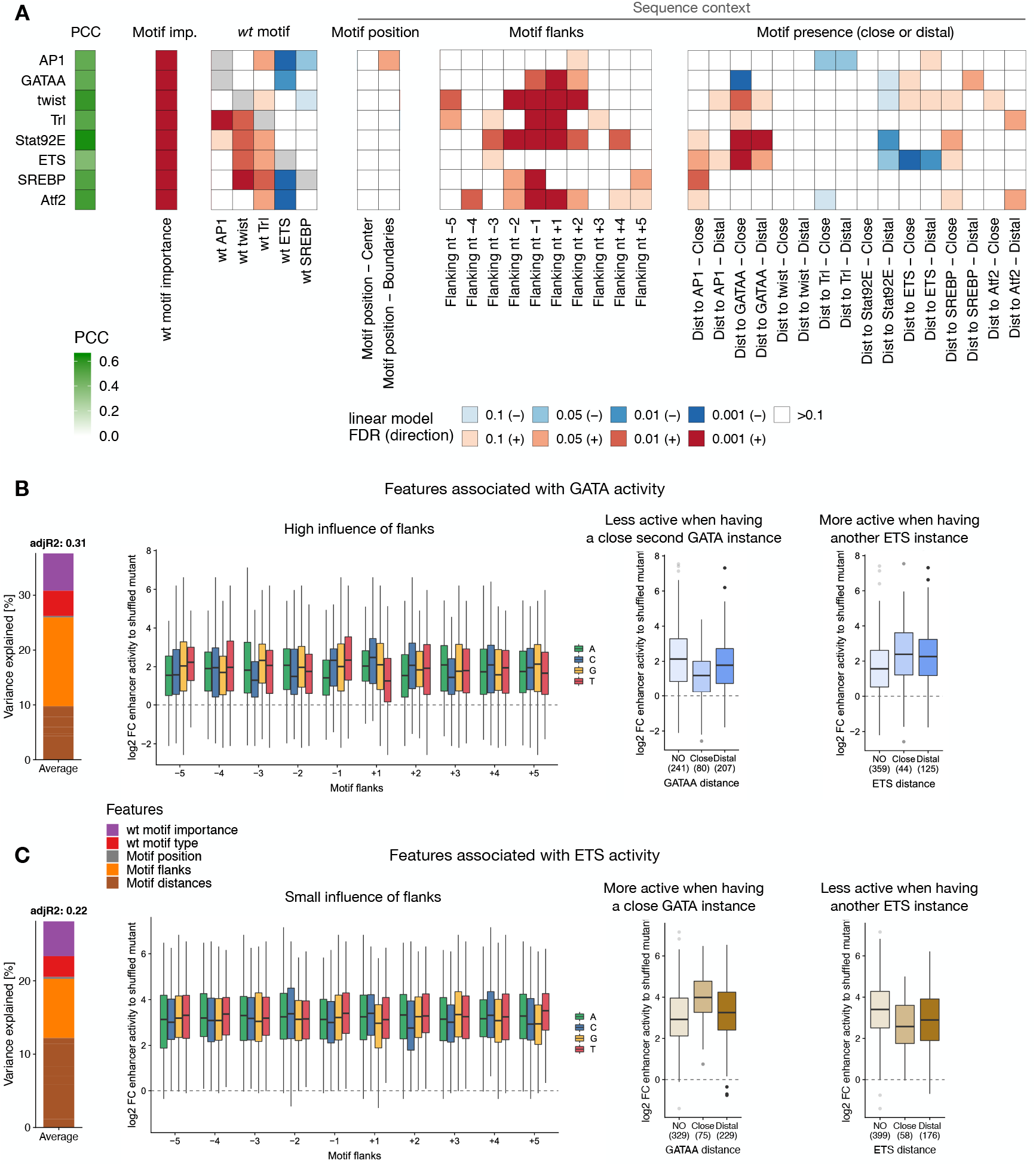
Characterization of preferred syntax features of each TF motif. **A)** Motif syntax rules modulate TF motif function. For each TF motif type (rows), a linear model was built to predict its activity across all enhancer positions, using as covariates the number of instances, the wildtype TF motif importance and identity, and sequence context features such as the position within the enhancer, the flanking nucleotides, and the presence at close or distal distances to all other TF motifs. The PCC between predicted and observed motif activities is shown with the green color scale on the left. Heatmap shows the contribution of each feature (columns) for each model, colored by the FDR-corrected p-value (red or blue scale depending on positive or negative association, respectively). **B,C)** Syntax features associated with GATA **(B)** or ETS **(C)** activity. Left: bar plot showing the variance explained by the different types of features (color legend) for each of the linear models. Middle-left: motif activity according to the different bases at each flanking position, colored by nucleotide identity. Middle-right and right: enhancer activity changes (log2 FC to mutated sequence) after pasting each TF motif in positions with no additional GATA (middle-right) or ETS (right) in the enhancer, or with additional GATA or ETS at close (<= 25 bp) or distal (>25 bp) distances. Number of instances are shown.

The TF motif type-specific models revealed how the sequence context rules differ between TF motif types, explaining the motif-specific enhancer position preferences. For example, GATA activity was strongly dependent on the flanking nucleotides and was modulated by the presence of a second GATA at close distance (negative interaction) or ETS motifs (positive interaction) (Fig 4B). We saw different associations for ETS activity, as expected by the different GATA and ETS activity profiles across all positions (Fig 3E). ETS activity was only mildly influenced by the flanking nucleotides but strongly by neighboring motifs: it was stronger close to GATA motifs and weaker in enhancers with another ETS motifs (Fig 4C). These sequence features, such as the negative GATA/GATA and the positive ETS/GATA interactions at close distances, were observed previously via computational models of wildtype S2 enhancer sequences (de Almeida et al. 2022).

In addition, analyzing the DeepSTARR predicted importance of each nucleotide when pasting different TF motifs at the same position revealed their interaction with the sequence context (Fig S16): GATA but not ETS activated the chr3L enhancer in a position with additional distal GATA motifs (note the increased weights for surrounding twist and AP-1 motifs when pasting GATA; Fig 4A,B), while ETS but not GATA activated the chrX enhancer in a position with a GATA motif at close distance (note the increased weights for the downstream GATA only when pasting ETS; Fig 4C,D), and both activated the chr2L enhancer that contains multiple surrounding twist motifs (Fig 4E,F). Together these results demonstrate how the sequence context (e.g. the flanking sequence, the presence and diversity of other motif types) modulates the function of TF motifs, constraining enhancer sequence flexibility.

### Enhancer sequence context modulates the function of human TF motifs

To test whether TF motifs also work differently in different enhancer sequence contexts in other species, we performed the systematic motif pasting experiment in human HCT116 cells for eight previously characterized human TF motifs (P53, AP-1, ETS, CREB1, MAF, EGR1, E2F1 and MECP2; see Methods; (de Almeida et al. 2022)). Pasting of the motifs into 1,354 important positions in 753 different HCT116 enhancers revealed that human TF motifs also have different intrinsic strengths and work differently in different enhancers and positions (Fig 5A, S17). P53 was the strongest motif and the only one that showed globally strong activity across all enhancer positions, suggesting little dependence on the enhancer context, as has been suggested before (Verfaillie et al. 2016). AP-1, the second strongest motif, was strongly dependent on the enhancer positions, with activities ranging more than 50-fold across enhancer contexts. This position-dependence was also observed for the other motifs, even though their overall activity was lower (Fig 5A).

**Figure 5.**
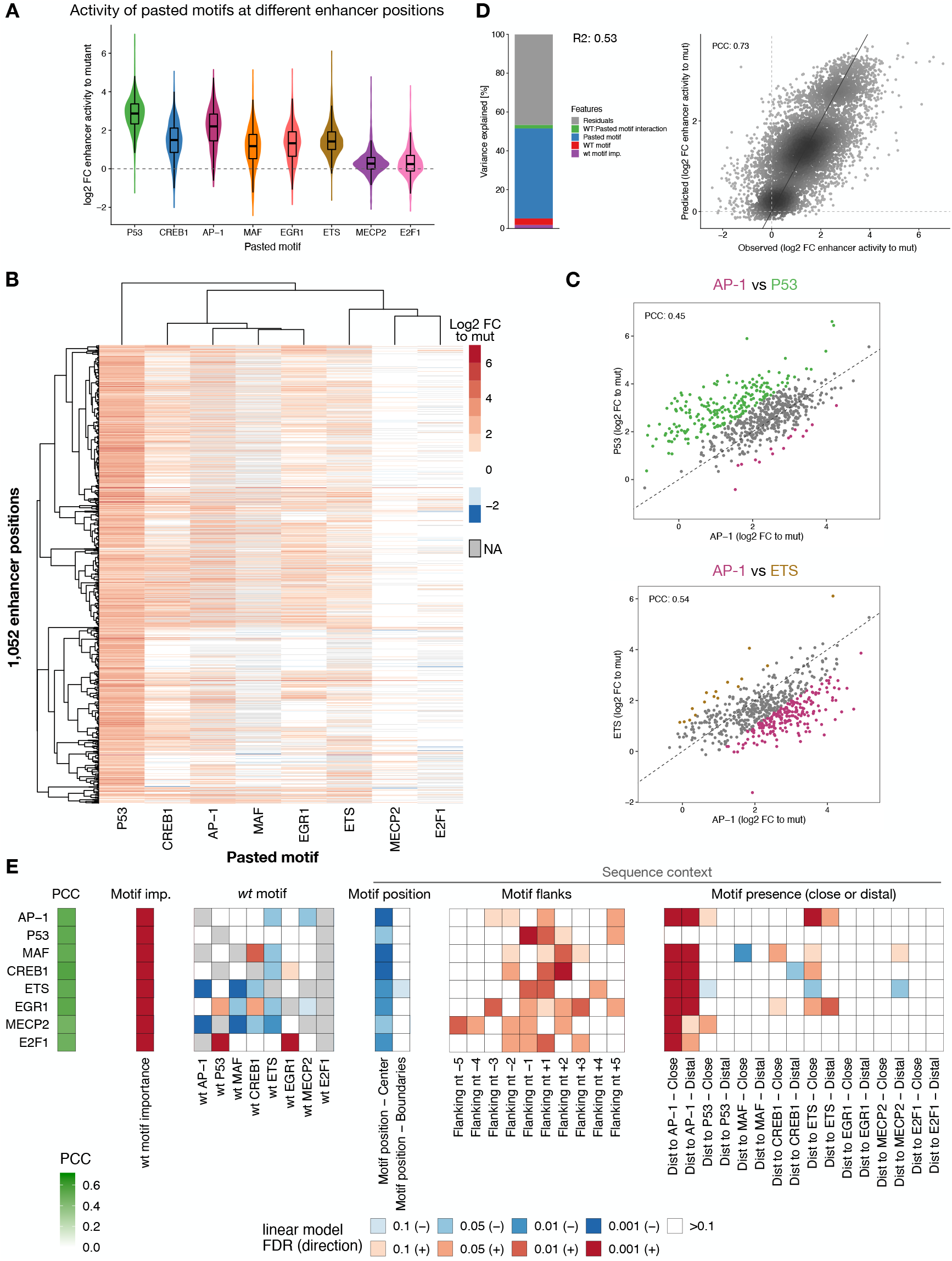
Human TF motifs require specific enhancer sequence contexts. **A)** Distribution of enhancer activity changes (log2 FC to mutated sequence) across all enhancer positions for each pasted TF motif. **B)** Heatmap of enhancer activity changes (log2 FC to mutated sequence) after pasting each of the eight selected human TF motifs in 1,052 enhancer positions (positions with data for at least six motifs). TF motifs and positions were clustered using hierarchical clustering and the activity is colored in shades of red (activating) and blue (repressing); missing values are colored in grey. **C)** Human TF motifs work differently at different enhancer positions. Comparison between enhancer activity changes (log2 FC to mutated sequence) after pasting AP-1 (x-axis) and P53 (top) or ETS (bottom) (y-axis), across all enhancer positions. Positions with stronger activity of each motif (>= 2-fold in respect to the other motif in the scatter plot) are colored (P53: green, AP-1: purple, ETS: brown). PCC: Pearson correlation coefficient. **D)** TF motif activity in function of wildtype and pasted motif identity. Left: Bar plot showing the amount of variance explained by the wildtype motif importance and identity, the pasted motif identity and the interaction between the wildtype and pasted motifs, using a linear model fit on all motif pasting results. Right: Scatter plots of predicted (linear model) vs. observed enhancer activity changes (log2 FC to mutated sequence) across all motif pasting experiments. Color reflects point density. **E)** Motif syntax rules modulate the function of human TF motifs. For each TF motif type (rows), we built a linear model to predict their activity across all enhancer positions, using as covariates the number of instances, the wildtype TF motif importance and identity, and sequence context features such as the position within the enhancer, the flanking nucleotides, and the presence at close or distal distances to all other TF motifs. The PCC between predicted and observed motif activities is shown with the green color scale on the left. Heatmap shows the contribution of each feature (columns) for each model, colored by the FDR-corrected p-value (red or blue scale depending on positive or negative association, respectively).

TF motifs preferred different enhancer contexts, with four groups of motifs showing characteristically different preferences: (1 – P53) strong activity in all positions; (2 – CREB1, AP-1, MAF, EGR1) and (3 – ETS) highly context-dependent activities; (4 – MECP2, E2F1) only active in few and highly specific enhancer positions (Fig 5B,C, S18). These distinct preferences were independent of the identity of the replaced motif (Fig 5D, S19) but correlated with sequence context features. Similar to *Drosophila* TF motifs, motif context features such as motif flanks and the presence and distance to other TF motifs were important to predict the activities of human motifs across the different enhancer positions (Fig S20). TF-specific linear models based on such syntax features were able to predict the motif activities across all positions (PCCs between 0.46 and 0.51; Fig S21) and revealed the context preferences of each TF motif (Fig 5E).

All motif activities were influenced by the flanking nucleotides, that explained on average 8.2% of the motif activities variance, while the presence of additional motifs and their distance explained 8.5% (Fig 5E, S21,22). As expected by the weak context-specificity of P53 (group 1, Fig 5A), its activity was independent of the presence and distance to other TF motifs (Fig 5E, S22A). All the other motifs preferred contexts with an additional AP-1 instance (Fig 5E). The AP-1 motif itself, as well as MAF, CREB1 and EGR1 (group 2), all preferred positions close to an ETS motif, concordant with previous studies showing direct protein–protein interactions between ETS and other TFs (Li et al. 2000; Burda et al. 2010), while the ETS motif (group 3) had a negative interaction with a second close ETS motif (Fig 5E), as also observed in *Drosophila* enhancers (Fig 4A). These findings are also concordant with the motif syntax rules found in a previous study (de Almeida et al. 2022). Altogether, this establishes that TF motifs require specific enhancer sequence contexts in species as divergent as fly and human, suggesting that this is a general principle of regulatory enhancer sequences.

## Discussion

In this study, we used two complementary strategies to explore the flexibility of enhancer sequences with regards to nucleotide and motif identity at specific enhancer positions as well as the position dependence of motif activity. Even though median enhancer activity drops significantly when randomizing an 8nt stretch at important enhancer positions, many sequence variants, including both variants of the wildtype motif as well as other TF motifs, can still achieve strong enhancer-activity. The diverse set of solutions found at each position shows that enhancers exhibit some degree of flexibility. However, as demonstrated by the fact that only a few hundred out of the 65,000 possible sequences work, the flexibility at any given position is constrained by the enhancer context and determined by syntax rules. Similarly, systematically pasting different motifs into hundreds of enhancer positions revealed that motif activity is strongly modulated by the enhancer sequence context, and neither is any motif able to functionally replace any other motif, nor are all motifs able to function at all positions. Therefore, constrained sequence flexibility and the modulation of motif function by the sequence context seem to be key features of enhancers.

The observation that both *Drosophila* and human TF motifs require specific enhancer sequence contexts suggests that this is a general principle of regulatory enhancer sequences. Even though motifs possess some intrinsic strength, this potential to activate transcription strongly depends on the enhancer context and follows certain syntax rules, including motif flanks, combinations and distances. While the motif flanking sequence can influence TF binding affinity via changes in DNA shape (Mathelier et al. 2016; Dror et al. 2015), inter-motif distances can impact the synergy between TFs at the level of DNA binding or after binding, such as cofactor recruitment and activation (Reiter et al. 2017). Although these rules are stricter for some TF motifs (e.g. GATA) and more relaxed for others (e.g. P53), motifs are not simply independent modules but interact with all enhancer features in a highly cooperative manner, which can modulate motif activity by more than 100-fold. This is an important result that supports a model where enhancer activity is encoded through a complex interdependence between motifs and context, rather than TF motifs acting independently and additively as the billboard model would suggest (Kulkarni and Arnosti 2003; Arnosti and Kulkarni 2005). While tissue- or cell-type-specificity can already be predicted by motif presence-absence patterns alone (Kvon et al. 2014; Janssens et al. 2022), the encoding of different enhancer strengths depends on more complex cis-regulatory syntax rules (de Almeida et al. 2022; Jindal and Farley 2021). Mutations in TF motifs and changes in the enhancer sequences can therefore only be understood in the context of these syntax features.

The motif syntax rules described here (Fig 4, 5E), such as the interaction between motifs and their distances, agree well with the ones learned by the DeepSTARR deep learning model trained on genome-wide enhancer activity data (de Almeida et al. 2022), showing that these rules are present and important in wildtype enhancer sequences. Indeed, DeepSTARR also predicted with good accuracy the activity of all randomized sequence variants and of motifs pasted in different enhancer contexts (Fig S23, S24). This supports the validity of computational models such as DeepSTARR and their use in *in-silico-like* experiments (e.g motif pasting experiments with a larger set of TF motifs across many more genomic positions) to improve our understanding of the regulatory information encoded in enhancer sequences and the impact of mutations.

Our study shows that enhancer sequences are flexible enough for enhancer strength to be achieved by a small yet diverse set of sequence variants, and that mutations in information-poor positions have little impact on the enhancer activity in a single cell type. This flexibility that allows many different sequences to achieve similar enhancer activities in a single cell type might be an important pre-requisite for the evolution of developmental enhancers that operate under many additional constraints, e.g. regarding the precise spatio-temporal control of enhancer activities. Given that the activity in a given cell can be achieved by many solutions, the specific solutions that fulfill additional requirements can be explored during evolution. Indeed, previous studies that have analyzed expression changes of enhancer mutations across different cell types *in vivo* have observed that the cell type-specific expression patterns of enhancers can change upon (minimal) sequence perturbations (Farley et al. 2015; Galupa et al. 2022; Fuqua et al. 2020). The fact that enhancer strength in any given cell type and specificity across cell types and developmental time are subject to different sequence constraints highlight the complexity of the regulatory code and the challenges faced when trying to dissect it. We expect that the combination of quantitative enhancer-sequence-to-function models in individual cell types with qualitative predictions of enhancer activities across cell types will over the next years provide unprecedented progress in our understanding of enhancer biology and our ability to read and write enhancer sequences.

## Methods

### UMI-STARR-seq

#### Cell culture and transfection

*Drosophila* Schneider 2 cells were grown in Schneider’s *Drosophila* Medium (Gibco; 21720-024) supplemented with 10% heat inactivated FBS (Sigma; F7524) at 27°C. Human HCT116 cells were cultured in DMEM (Gibco; 52100-047) supplemented with 10% heat inactivated FBS (Sigma; F7524) and 2mM L-Glutamine (Sigma; G7513) at 37°C in a 5% C0_2_-enriched atmosphere. Both cell types were passaged every 2-3 days.

We used the MaxCyte-STX electroporation system for all library transfections. S2 cells were collected at 300 x g for 5min and washed once in 1:1 Schneider’s Drosophila Medium and MaxCyte electroporation buffer (EPB-1). 50 x 10^6^ cells were transfected with 5µg of DNA using the “Optimization 1” protocol, recovered for 30min at 27°C and resuspended in 10mL S2 Medium with 10% FBS. HCT116 cells were collected at 200 x g for 5min and washed once in MaxCyte electroporation buffer (EPB-1). Cells were electroporated at a density of 1 x 10^7^ cells per 100µL and 20µg of DNA using the preset “HCT116” program, recovered for 20min at 37 °C and resuspended in 10mL DMEM 10% FBS and 2mM L-Glutamine.

Each replicate for a STARR-seq screen was transfected in 2 OC400 cuvettes with a total of 400 x 10^6^.

#### UMI-STARR-seq experiments

##### Library cloning

Random 8nt variant libraries were generated using a PCR approach with degenerate oligonucleotides. Forward primers (primers see Supplementary Table S1) were designed to anneal directly downstream of the enhancer position of interested followed by 8 degenerate bp (creating 65,536 variants) and another 20 bp complementary stretch. Reverse primers were complementary to the 20 bp 5’ of the degenerate stretch. The STARR-seq vector containing the wildtype enhancer of interest (either *ced-6* or *ZnT63C*) was used as a template for the PCR. The PCR was run across the whole STARR-seq plasmid, followed by DpnI digest and a Gibson reaction that re-circularizes the plasmid. Libraries were grown in 2l LB-Amp (final ampicillin concentration 100µg/mL). Variant libraries of the same enhancer i.e. *ced-6* enhancer pos110, pos182, pos230, pos241 and *ZnT63C* enhancer pos142, pos180, pos210 were pooled to equimolar ratio, together with another synthetic oligo library containing wt enhancer sequences and negative regions.

*Drosophila* and human oligo libraries were synthesized by Twist Bioscience including the 249 bp enhancer sequence and adaptors for library cloning. *Drosophila* library fragments were amplified (primers see Supplementary Table S1) and cloned into *Drosophila* STARR-seq vectors containing the DSCP core-promoters using Gibson cloning (New England BioLabs; E2611S). The oligo library for human STARR-seq screens was amplified (primers see Supplementary Table S1) and cloned into the human STARR-seq plasmid with the ORI in place of the core promoter (Muerdter et al. 2018). Libraries were grown in 2l LB-Amp (final ampicillin concentration 100µg/mL).

All libraries were purified with Qiagen Plasmid *Plus* Giga Kit (cat. no. 12991).

##### Drosophila S2 cells

UMI-STARR-seq was performed as described previously (Arnold et al. 2013; Neumayr et al. 2019). In brief, we transfected 400 × 10^∧^6 S2 cells total per replicate with 20 μg of the input library (see libraries above). After 24 hr incubation, poly-A RNA was isolated and processed as described before (Neumayr et al. 2019). Briefly: after reverse transcription and second strand synthesis a unique molecular identifier (UMI) was added to each transcript, allowing the counting of individual RNA molecules. This is followed by two nested PCR steps, each with primers that are specific to the reporter transcripts such that STARR-seq does not detect endogenous cellular RNAs.

##### Human HCT116 cells

UMI-STARR-seq was performed as described previously (Arnold et al. 2013; Muerdter et al. 2018; Neumayr et al. 2019). Screening libraries were generated from synthesized oligo pools by Twist Bioscience (see above). We transfected 80 × 10^∧^6 HCT116 cells total per replicate with 160 μg of the input library. After 6 hr incubation, poly-A RNA was isolated and further processed as described before (Neumayr et al. 2019).

##### Illumina sequencing

Next-generation sequencing was performed at the VBCF NGS facility on an Illumina NextSeq 550 or NovaSeq SP platform, following manufacturer’s protocol. Random variants UMI-STARR-seq and Twist-oligo library screens were sequenced as paired-end 150 cycle runs, using standard Illumina i5 indexes as well as unique molecular identifiers (UMIs) at the i7 index. Deep sequencing base-calling was performed with CASAVA (v.1.9.1).

#### Random variants UMI-STARR-seq data analysis

Dedicated bowtie indices were created for each enhancer position’s N_8_ library and combined with an oligo library of thousands of wildtype enhancers and negative sequences (de Almeida et al. 2022) for normalization, all 249 bp-long sequences. UMI-STARR-seq RNA and DNA input reads (paired-end 150 bp) were mapped to these dedicated bowtie indices using Bowtie v.1.2.2 (Langmead et al. 2009). Since the N_8_ variants were all positioned in the last 150 nt of each enhancer, we allowed for flexible mapping in the beginning of the fragments to increase the number of mapped reads while keeping high sensitivity for the different enhancer variants. Specifically, we trimmed the forward reads to 36 bp and mapped them to the indices allowing for 3 mismatches; the full 150 bp-long reverse reads were mapped with no mismatches, to identify all sequence variants; paired-end reads with the correct position, length and strand were kept. This mapping strategy was used for both DNA and RNA reads. For paired-end DNA and RNA reads that mapped to the same variant, we collapsed those that have identical UMIs (10 bp, allowing one mismatch) to ensure the counting of unique molecules (Supplementary Table 2).

We excluded oligos with less than 5 reads in any of the input replicates and less than 1 read in any of the RNA replicates. The enhancer activity of each sequence in each screen was calculated as the log2 fold-change over input, using all replicates, with DESeq2 (Love et al. 2014). We used the counts of wildtype negative regions in each library as scaling factors between samples.

#### Oligo library UMI-STARR-seq data analysis

As described previously (de Almeida et al. 2022), oligo library UMI-STARR-seq RNA and DNA input reads (paired-end 150 bp) were mapped to a reference containing the 249 bp-long sequences from the fragments present in the *Drosophila* (dm3) or human (hg19) libraries using Bowtie v.1.2.2 (Langmead et al. 2009). For each library we demultiplexed reads by the i5 and i7 indexes and oligo identity. Mapping reads with the correct length, strand and with no mismatches (to identify all sequence variants) were kept. Both DNA and RNA reads were collapsed by UMIs (10 bp) as above (Supplementary Table 2).

We excluded oligos with less than 10 reads in any of the input replicates and added one read pseudocount to oligos with zero RNA counts. The enhancer activity of each oligo in each screen was calculated as the log2 fold-change over input, using all replicates, with DESeq2 (Love et al. 2014). We used the counts of wildtype negative regions in each library as scaling factors between samples.

### Analyses of random variants at different enhancer positions

#### Independent motif mutations

Two strong S2 developmental enhancers with different TF motif compositions were selected to test a diversity of random 8 nt variants in different positions: *ced-6* (chr2R:5326628-5326876) and *ZnT63C* (chr3L:3310914-3311162) enhancers. Experimental mutations of GATA, AP-1 and twist motifs in these enhancers were performed in a previous study (Fig S4F; (de Almeida et al. 2022)) and used here to select important enhancer positions.

#### Enhancer random variants libraries and UMI-STARR-seq

We selected five positions important for the activity of the two enhancers (*ced-6* pos110 and pos241; *ZnT63C* pos142, pos180, pos210) and two non-important positions of the *ced-6* enhancer (pos182 and pos230). At each position, we experimentally replaced the respective 8nt stretch of the enhancer with randomized nucleotides (N_8_), creating 65,535 enhancer variants in addition to the wildtype sequence per position. For each enhancer, we pooled the libraries of the different positions and combined them with an oligo library of thousands of wildtype enhancers and negative sequences (de Almeida et al. 2022) for normalization. UMI-STARR-seq using the *ced-6* or *ZnT63C* pooled libraries was performed (“UMI-STARR-seq experiments”) and analyzed (“Random variants UMI-STARR-seq data analysis”) as described above (Supplementary Table 3). We performed two independent replicates per enhancer pooled library screen (Pearson correlation coefficient (PCC)=0.85-0.91; Fig S4A-E).

To be able to compare the activity of variants and motifs between enhancer positions, we next scaled the enhancer activity of all variants per position (z-scores). This allows to measure the change in activity of a given variant over the average of all variants, correcting for the importance of the different enhancer positions tested.

#### Comparison between pooled libraries using common oligos

The respective wildtype enhancer sequence was overrepresented in each N_8_ library input since it was used as the template for the PCR cloning (Fig S4A,B). We compared the activities of the *ced-6* and *ZnT63C* enhancer sequences and all other wildtype enhancers and negative sequences present in both *ced-6* and *ZnT63C* pooled libraries (Fig S4C). The activities of the common sequences were similar between both screens, except for the *ZnT63C* enhancer whose activity was underestimated in the *ZnT63C* pooled library, likely due to the technical overrepresentation in the input. We therefore selected another enhancer with the same activity as the *ZnT63C* enhancer (chrX:9273894-9274142) to be used as the reference wildtype activity for the *ZnT63C* enhancer variants (Fig S4C, 2B).

#### Diversity of top active variants and *de novo* motif discovery

The most-active 8nt variants of each screen (1, 2, 5, 10, 50, 100 and 1,000) were retrieved and consolidated into position probability matrices based on the nucleotide frequencies at each position (Fig 1C, S5B). Logos were visualized using the *ggseqlogo* function from R package *ggseqlogo* (v.0.1; (Omar Wagih 2017)). The same was done after randomly sorting the variants of each screen for comparison. The information content of the top sequences at each position was calculated as described in https://bioconductor.org/packages/release/bioc/vignettes/universalmotif/inst/doc/IntroductionToSequenceMotifs.pdf (Schneider and Stephens 1990; Schneider et al. 1986) (Fig 1D, S5C).

The top 100 and 1,000 or bottom 1,000 variants (8nt +/-4nt flanks) of each screen were used for *de novo* motif discovery analyses using HOMER, taking all detected variants of the respective screen as background (Fig S2, S6). HOMER (v4.10.4; (Heinz et al. 2010)) was run with the findMotifs.pl command and the arguments *fly-len 6,7,8*.

#### Activity of TF motifs created by sequence variants

To robustly assess the activity of a given TF motif, we retrieved the activity of all 16nt variants (8nt +/-4nt flanks) creating each motif by string-matching. The main motifs used were: GATA – GATAAG, AP-1 – TGA.TCA, SREBP – TCACGCGA, twist – CATCTG, CREB/ATF – TCATCA, STAT – TTCC.GGA, Trl – GAGAGA, ETS – CCGGAA, Dref – ATCGAT, ttk – AGGATAA, ZEB1 – CAGGTG, lola – GGAGTT (format: TF motif – string). For a more systematic comparison across all TF motif types, we matched variants to the optimal string from each TF motif PWM model in a motif database (Fig S8A; (de Almeida et al. 2022)). The average activity across variants was defined as the motifs’ intrinsic strength. These activities were used in Fig 1E, S3A, 2E,D, S8, S9.

To find how many active variants are explained by the creation of known motifs enriched in S2 developmental enhancers (from (de Almeida et al. 2022)), we performed PWM-based motif scanning of those candidate motifs onto variants (8nt +/-4 flanks) (Fig 1F, S3B). We used the *matchMotifs* function from R package *motifmatchr* (v.1.4.0; genome = “BSgenome.Dmelanogaster.UCSC.dm3”, bg=“genome” (Schep 2021)) with p-value cutoffs 1e^-04^ and 1e^-05^.

#### Comparison of random variants activity across enhancer positions

We compared the activity of all 8nt random variants across enhancer positions using their z-score scaled activity (Fig 2C, S7; Supplementary Table 3). We calculated pairwise PCCs between the different libraries, performed hierarchical clustering (“complete” method) using the correlation values as similarities, and displayed heatmaps using the *pheatmap* R package (v.1.0.12; (Kolde 2019)). To reduce the impact of the flanking sequence of each position when comparing the activity of variants between them, we repeated the same after consolidating the 8nt into shorter variants by taking the centered sequence and averaging the activity across variants with different flanking nucleotides.

### Analyses of motif pasting screens in *Drosophila* and human enhancers

#### Oligo library design

##### *Drosophila* motif pasting library

We selected 1,172 motif positions (among 728 enhancers) that are required for the activity of the respective enhancers, assessed by experimental mutagenesis in a previous study (de Almeida et al. 2022). These wildtype positions cover different contexts and TF motifs: GATA, AP-1, twist, Trl, ETS and SREBP. We next designed sequences of enhancer variants where we pasted a mutant sequence or the optimal sequence of eight TF motifs (GATA, AP-1, twist, Trl, ETS, SREBP, Stat92E and Atf2; one at a time; sequences in Supplementary Table 4) in each of these positions (Fig 3A). Importantly, we pasted an extended optimal sequence of each TF motif to reduce the influence of flanking nucleotides and different motif affinities and focus on differences due to the enhancer context. This library (Supplementary Table 5) was synthetized and pooled with a previous library containing the wildtype enhancer sequences (de Almeida et al. 2022) to be screened together.

##### Human motif pasting library

Similar to the *Drosophila* library, we selected 1,456 motif positions important for the activity of 808 enhancers, assessed by experimental mutagenesis in a previous study (de Almeida et al. 2022). These wildtype positions cover different contexts and TF motifs: AP1, ETS, E2F1, EGR1, MAF, MECP2, CREB1, P53. We next designed sequences of enhancer variants where we pasted a mutant sequence or the optimal sequence of the same eight TF motifs (AP1, ETS, E2F1, EGR1, MAF, MECP2, CREB1, P53; one at a time; sequences in Supplementary Table 4) in each of these positions. Importantly, we pasted an extended optimal sequence of each TF motif to reduce the influence of flanking nucleotides and different motif affinities and focus on differences due to the enhancer context. This library (Supplementary Table 6) was synthetized and pooled with a previous library containing the wildtype enhancer sequences (de Almeida et al. 2022) to be screened together.

#### Oligo library synthesis and UMI-STARR-seq

The *Drosophila* and human enhancers’ oligo libraries contained each sequences for the wildtype enhancers and enhancers with mutant variants or motifs pasted at the selected positions (Supplementary Table 5 and 6, respectively). All sequences were designed using the dm3 and hg19 genome versions, respectively. The enhancer sequences spanned 249 bp total, flanked by the Illumina i5 (25 bp; 5′-TCCCTACACGACGCTCTTCCGATCT) and i7 (26 bp; 5′ AGATCGGAAGAGCACACGTCTGAACT) adaptor sequences upstream and downstream, respectively, serving as constant linkers for amplification and cloning. The resulting 300-mer oligonucleotide *Drosophila* and human libraries were synthesized by Twist Bioscience. UMI-STARR-seq using these oligo libraries was performed (“UMI-STARR-seq experiments”) and analyzed (“Oligo library UMI-STARR-seq data analysis”) as described above (Supplementary Table 5 and 6). We performed three independent replicates for *Drosophila* (correlation PCC=0.95-0.98; Fig S10A,B) and human (PCC=0.96-0.98; Fig S17A,B) screens.

#### Quantification of motif activity at different enhancer positions

We used our enhancer activity measures of the wildtype and mutated sequences to stringently select important enhancer positions for further analyses: positions where mutation reduced the activity by at least 2-fold (Fig S10C, S17C). These resulted in 763 important positions distributed among 496 *Drosophila* enhancers and 1,354 positions distributed among 753 human enhancers. This was important to select positions where we could reliably measure the increase in enhancer activity after pasting each TF motif – quantified as the log2 fold-change activity over the mutated enhancer (Fig 3B, 5A). Variability of activity of each motif across enhancer positions was quantified using the coefficient of variation (ratio of the standard deviation to the mean; Fig S10D).

We compared the activity of motifs across enhancer positions by pairwise PCCs and performed hierarchical clustering (“complete” method) using the correlation values as similarities. Heatmaps were displayed using the *pheatmap* R package (v.1.0.12; (Kolde 2019)) (Fig 3D, S11A, 5B, S18A).

#### Importance of the wildtype motif

We fitted motif activity values (log2 fold-change enhancer activity after motif pasting) with linear models using the wildtype TF motif identity and importance (log2 fold-change activity between wildtype and motif-mutant sequence), the pasted motif identity, and the interaction between the wildtype and pasted motifs as covariates, using the *lm* function (v.3.5.1; (R Core Team 2020)). Variance explained by each covariate was calculated with one-way ANOVAs of the respective models (Fig S12B, 5D, S19B).

#### Difference between pairs of positions in the same or different enhancers

*Drosophila* enhancers with two positions tested in our assay were selected and the fold-change in motif activity between pairs of positions in the same enhancer was compared with the fold-change between pairs of positions in different enhancers (matched by similar position-mutant baseline activities). For each pasted TF motif, significant differences were assessed through a two-sided Wilcoxon signed rank test followed by FDR multiple testing correction (Fig S13).

#### Prediction of motif activities using motif syntax features

##### Motif syntax features

To test how motif activities depend on motif syntax features we extracted the following features per tested enhancer position: the position relative to the enhancer center (center:-/+ 25 bp, flanks: -/+25:75 bp, boundaries: -/+75:125 bp), the position flanking nucleotides (5 bp on each side), and the presence and distance to other TF motifs (close: <= 25 bp; distal: >25 bp; between motif centers).

Instances of each TF motif type were mapped across all enhancers using their annotated PWM models (Supplementary Table 3) and the *matchMotifs* function from R package *motifmatchr* (v.1.4.0; (Schep 2021)) with the following parameters: genome = “BSgenome.Dmelanogaster.UCSC.dm3”, p.cutoff = 5e-04, bg=“genome”. Overlapping instances (minimum 50%) for the same TF motif were collapsed and counted only once.

##### Random forest models

We used a 10-fold cross-validation scheme to train random forest models to predict *Drosophila* or human motif pasting activities (log2 fold-change to mutant) using as features the wildtype TF motif identity and importance (log2 fold-change activity between wildtype and motif-mutant sequence) and the pasted motif identity, together or not with additional syntax features (described above). All models were built using the *Caret* R package (v. 6.0-80; (Kuhn 2018)) and feature importance was calculated using its *varImp* function. Predictions for each held-out test sets were used to compare with the observed motif activities and assess model performance (Fig S14, S20).

##### Linear model with motif syntax rules to predict motif activities

For each TF motif type, we built a multiple linear regression model to predict its activity (log2 fold-change to mutant) across different enhancer positions using as covariates the wildtype TF motif identity and importance (log2 fold-change activity between wildtype and motif-mutant sequence) together with additional syntax features (described above). All models were built using the Caret R package (v. 6.0-80; (Kuhn 2018)) and 10-fold cross-validation. Predictions for each held-out test sets were used to compare with the observed log2 fold-changes and assess model performance (Fig S15, S21).

The linear model coefficients and respective FDR-corrected p-values were used as metrics of importance for each feature, using the red or blue scale depending on positive or negative associations (Fig 4A, 5E). For flanking positions, we used always red because the direction of the association is not relevant. In addition, we calculated the percentage of variance explained by each covariate in the linear models built for each TF motif with one-way ANOVAs. For each TF motif, we generated 100 different models, randomizing the order of the covariates (since the variance explained depends on the order of covariates entered), quantified the percentage of variance explained of each covariate as its sum of squares divided by the total sum of squares, and used the average value across all 100 models as the final variance explained per covariate (Fig S15, S21).

### DeepSTARR predictions

#### Nucleotide contribution scores

Nucleotide contribution scores for wildtype enhancers or enhancer variants (Fig 2A, S5A, S11C,D, S16) were calculated as described previously (de Almeida et al. 2022), using DeepExplainer (the DeepSHAP implementation of DeepLIFT, see refs. (Shrikumar et al. 2017; Lundberg and Lee 2017; Lundberg et al. 2020); update from https://github.com/AvantiShri/shap/blob/master/shap/explainers/deep/deep_tf.py) and visualized using the *ggseqlogo* function from R package *ggseqlogo* (v.0.1; (Omar Wagih 2017)).

#### DeepSTARR predictions of enhancer sequence changes

DeepSTARR (https://github.com/bernardo-de-almeida/DeepSTARR, (de Almeida et al. 2022)) was used to predict the enhancer activity of N_8_ variants in enhancers (Fig S23) or the log2 fold-change enhancer activity of motif pasting sequences (Fig S24).

#### Statistics and data visualization

All statistical calculations and graphical displays have been performed in R statistical computing environment (v.3.5.1; (R Core Team 2020)) and using the R package *ggplot2* (Wickham 2016). In all box plots, the central line denotes the median, the box encompasses 25th to 75th percentile (interquartile range) and the whiskers extend to 1.5× interquartile range.

## Data and code availability

All raw and processed sequencing data generated in this study have been submitted to the NCBI Gene Expression Omnibus (GEO; https://www.ncbi.nlm.nih.gov/geo/) under accession number GSE211659 or Zenodo at https://doi.org/10.5281/zenodo.7010528. Code used to process the UMI-STARR-seq data as well as to reproduce all analyses, results and figures is available at https://github.com/bernardo-de-almeida/Variant_STARRseq.

## Acknowledgements

We thank V. Loubiere and T. Pachano (IMP) for comments on the manuscript and all members of the Stark group for discussions. Deep sequencing was performed at the Vienna Biocenter Core Facilities GmbH. Franziska Reiter is a recipient of a DOC Fellowship of the Austrian Academy of Sciences at the Research Institute of Molecular Pathology. Research in the Stark group is supported by the Austrian Science Fund (FWF). Basic research at the IMP is supported by Boehringer Ingelheim GmbH and the Austrian Research Promotion Agency (FFG). For the purpose of Open Access, the author has applied a CC-BY-NC-ND 4.0 International license to any Author Accepted Manuscript (AAM) version arising from this submission.

## Author Contributions

F.R., B.P.d.A. and A.S. conceived the project. F.R. performed all experiments. B.P.d.A. performed all computational analyses. F.R., B.P.d.A. and A.S. interpreted the data and wrote the manuscript. A.S. supervised the project.

## Supplementary figures

**Supplementary Figure 1.**
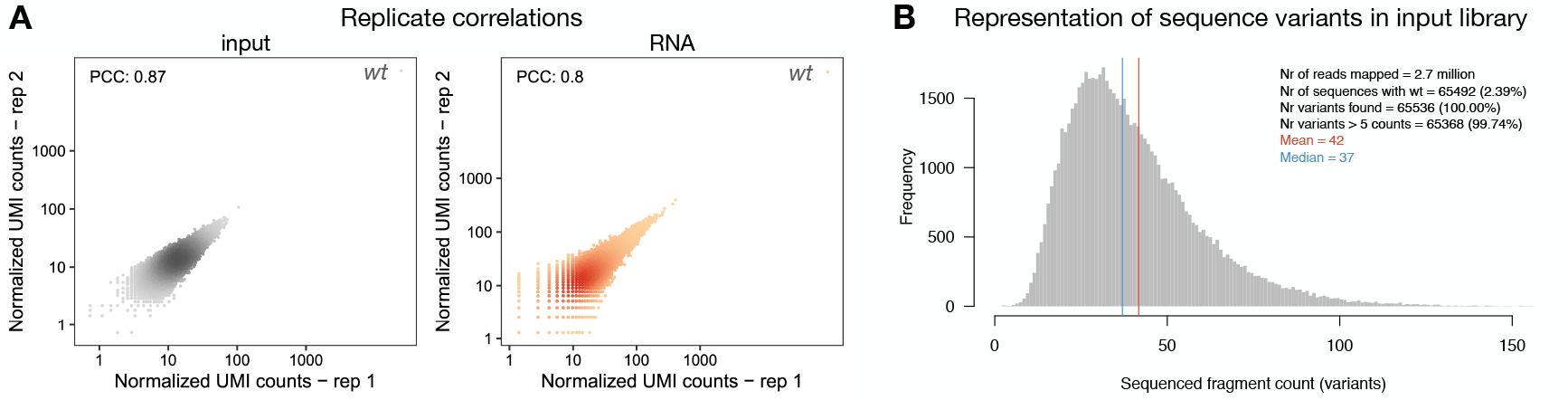
STARR-seq comprehensively assesses the activity of random variants in a specific region of the enhancer. **A)** Pairwise comparisons of normalized STARR-seq input (left) and RNA (right) UMI read counts between two independent biological replicates across all sequence variants tested in the GATA position (pos241) in the *ced-6* enhancer. Color reflects point density. The PCC is denoted for each comparison. Note the overrepresentation of the wildtype sequence both in the input and RNA libraries (top right corner), since it was used as the template for the PCR cloning (see Methods). **B)** Representation of sequence variants in STARR-seq input library. Frequency of variants covered by different number of UMI read counts. Number of sequences matching to wildtype and the number of variants recovered are shown, together with the mean and median counts sequenced per variant.

**Supplementary Figure 2.**
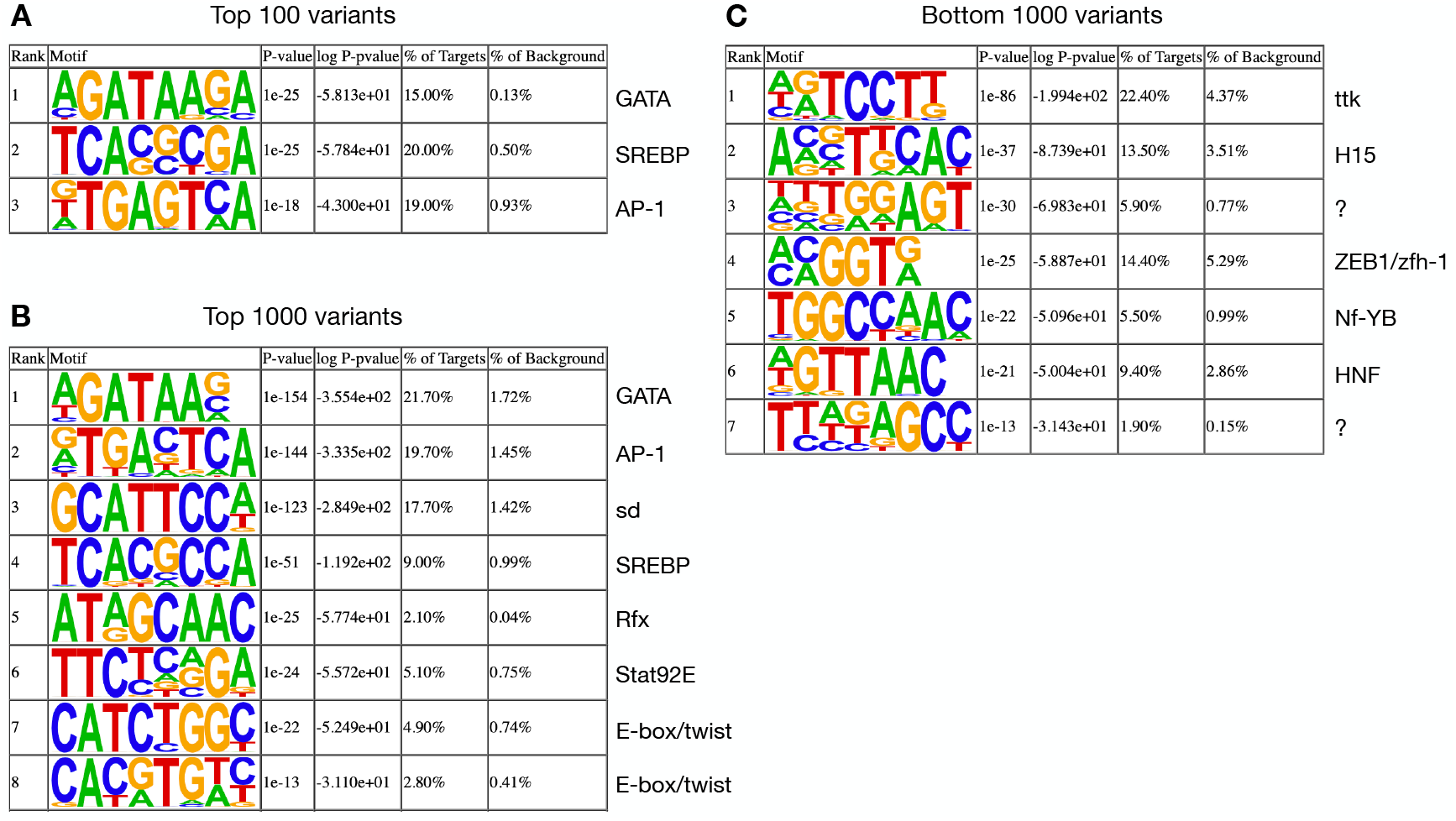
*De novo* motif discovery with Homer of top and bottom variants at the GATA position (pos241) in the *ced-6* enhancer. TF motifs found *de novo* (Homer) within the top 100 **(A)**, top 1,000 **(B)** or bottom 1,000 **(C)** variants. Motifs logo, statistics and predicted TF are shown.

**Supplementary Figure 3.**
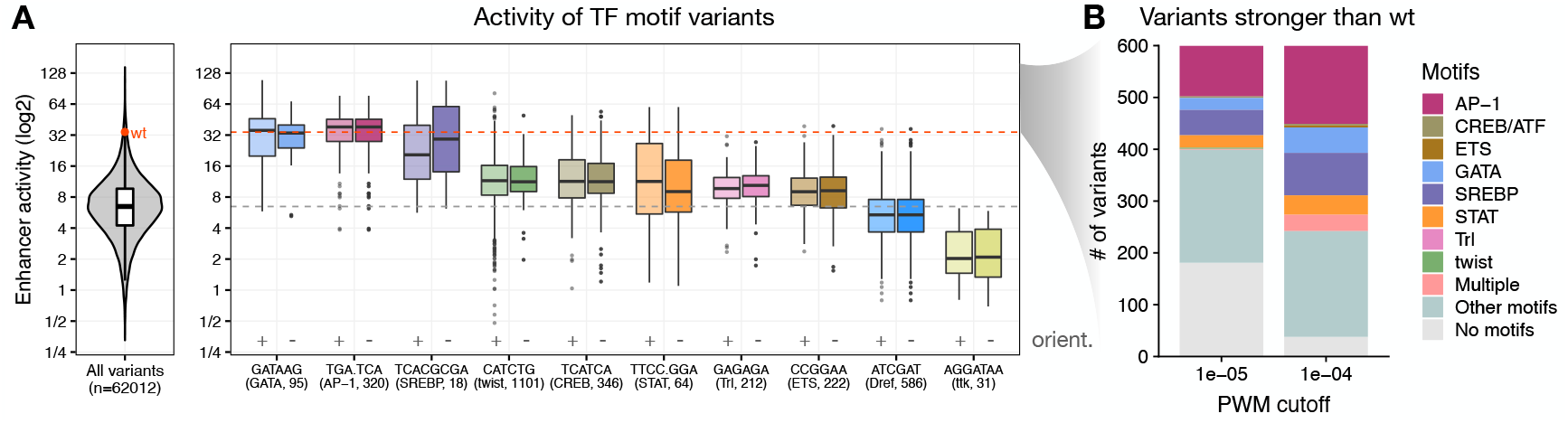
Activity of variants creating different TF motif types at the GATA position (pos241) in the *ced-6* enhancer. **A)** Distribution of enhancer activity for all 62,012 enhancer variants (left) or variants creating each TF motif in either orientation (right; positive and negative orientation are shown in grey). The activity of the wildtype sequence (wt, red dot and dashed line) or median of all variants (grey dashed line) are highlighted. The string of each TF motif used for the motif matching and the number of variants matching to each motif are described in the x-axis in the format “motif string (TF motif name, number of variants)”. **B)** Number of variants among the 600 stronger than wildtype that match to motifs enriched in S2 developmental enhancers, using two different PWM p-value cutoffs (1e^-05^ and 1e^-04^).

**Supplementary Figure 4.**
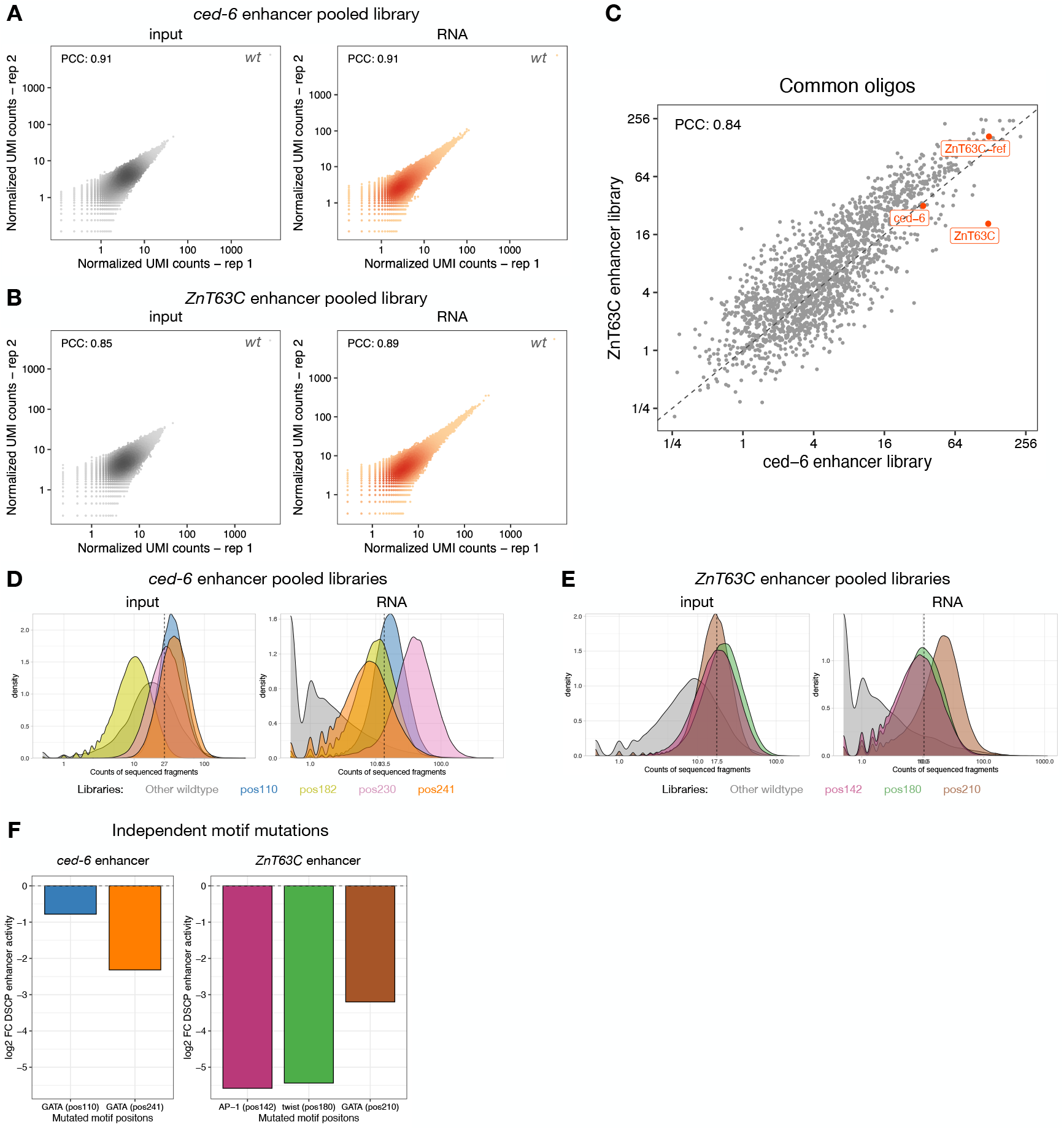
STARR-seq screens with random variants in seven positions of two different enhancers. **A,B)** Pairwise comparisons of normalized STARR-seq input (left) and RNA (right) UMI read counts between two independent biological replicates across all sequence variants tested in positions of the *ced-6* **(A)** or *ZnT63C* **(B)** enhancer. Color reflects point density. The PCC is denoted for each comparison. Note the overrepresentation of the wildtype sequence both in the input and RNA libraries (top right corner), since it was used as the template for the PCR cloning (see Methods). **C)** Comparison of enhancer activity between the two different enhancer pooled libraries for the common oligos (a library of wildtype enhancer or negative sequences; see Methods). The PCC is shown. The respective wildtype enhancers are highlighted. Given the underestimation of the activity of the *ZnT63C* wildtype sequence in its pooled library, we used as reference wildtype activity the activity of another enhancer with similar activity that was conserved in both libraries (see Methods). **D,E)** Representation of sequence variants from each individual library (a library of wildtype enhancer and negative sequences, grey, or libraries with random variants in each enhancer position, different colors) in STARR-seq input and RNA pooled libraries of the *ced-6* **(D)** or *ZnT63C* **(E)** enhancer. The mean counts sequenced per variant is shown per pooled library with a dashed line. **F)** Importance of each motif position selected in the *ced-6* (Left) or *ZnT63C* (Right) enhancer as judged by the impact of their individual mutation in enhancer activity (log2 fold-change). Data retrieved from *de Almeida et al., 2022*.

**Supplementary Figure 5.**
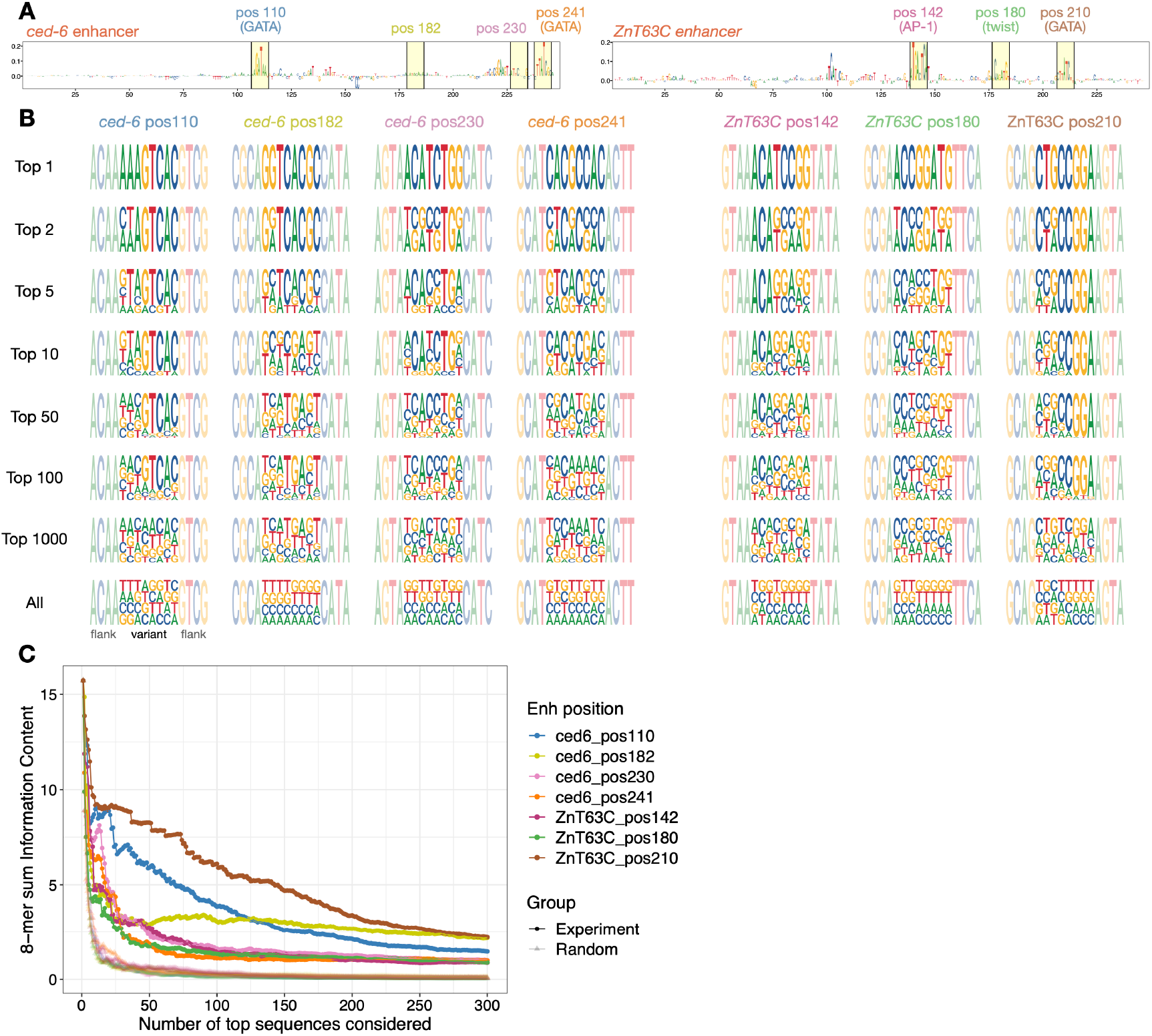
Top active variants at each enhancer position are highly diverse. **A)** DeepSTARR-predicted nucleotide contribution scores for the *ced-6* (left) and *ZnT63C* (right) selected enhancer sequences. Selected 8nt motif positions and non-important control positions are highlighted in yellow with the respective numerical position, TF motif identity and different colors. **B)** Strong sequence variants are highly diverse. Logos with nucleotide frequency of the most-active variants in STARR-seq (1, 2, 5, 10, 50, 100, 1,000 and all) at each enhancer position (colored as in (A)). **C)** Sum of information content within the most-active 8-mers in STARR-seq (colored as in (A)) compared with the same after randomly sorting the variants (grey) for each enhancer position, considering different number of top sequences.

**Supplementary Figure 6.**
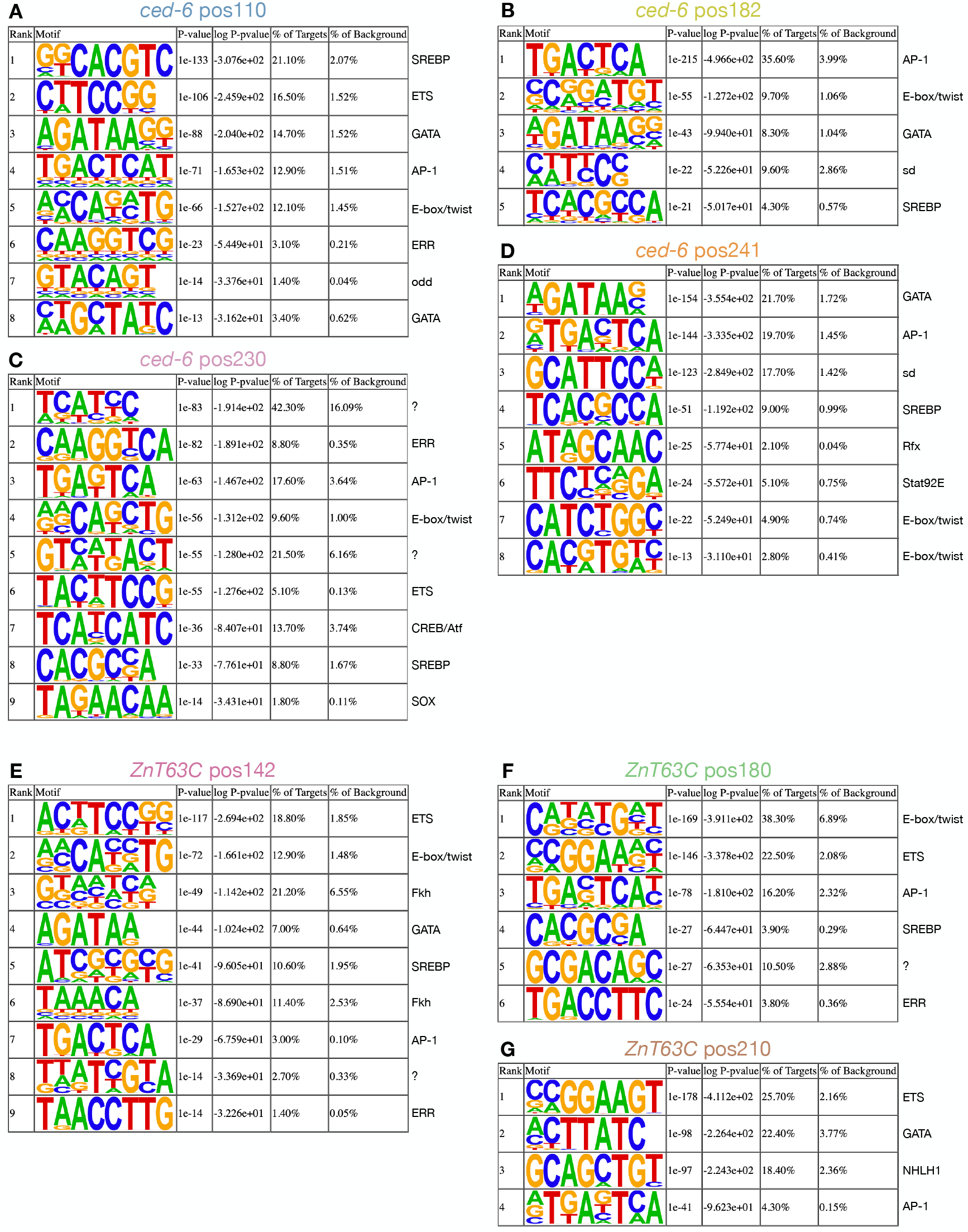
*De novo* motif discovery with Homer of the top 1000 variants at the different enhancer positions. TF motifs found *de novo* (Homer) within the top 1,000 variants at each enhancer position. Motifs logo, statistics and predicted TF are shown.

**Supplementary Figure 7.**
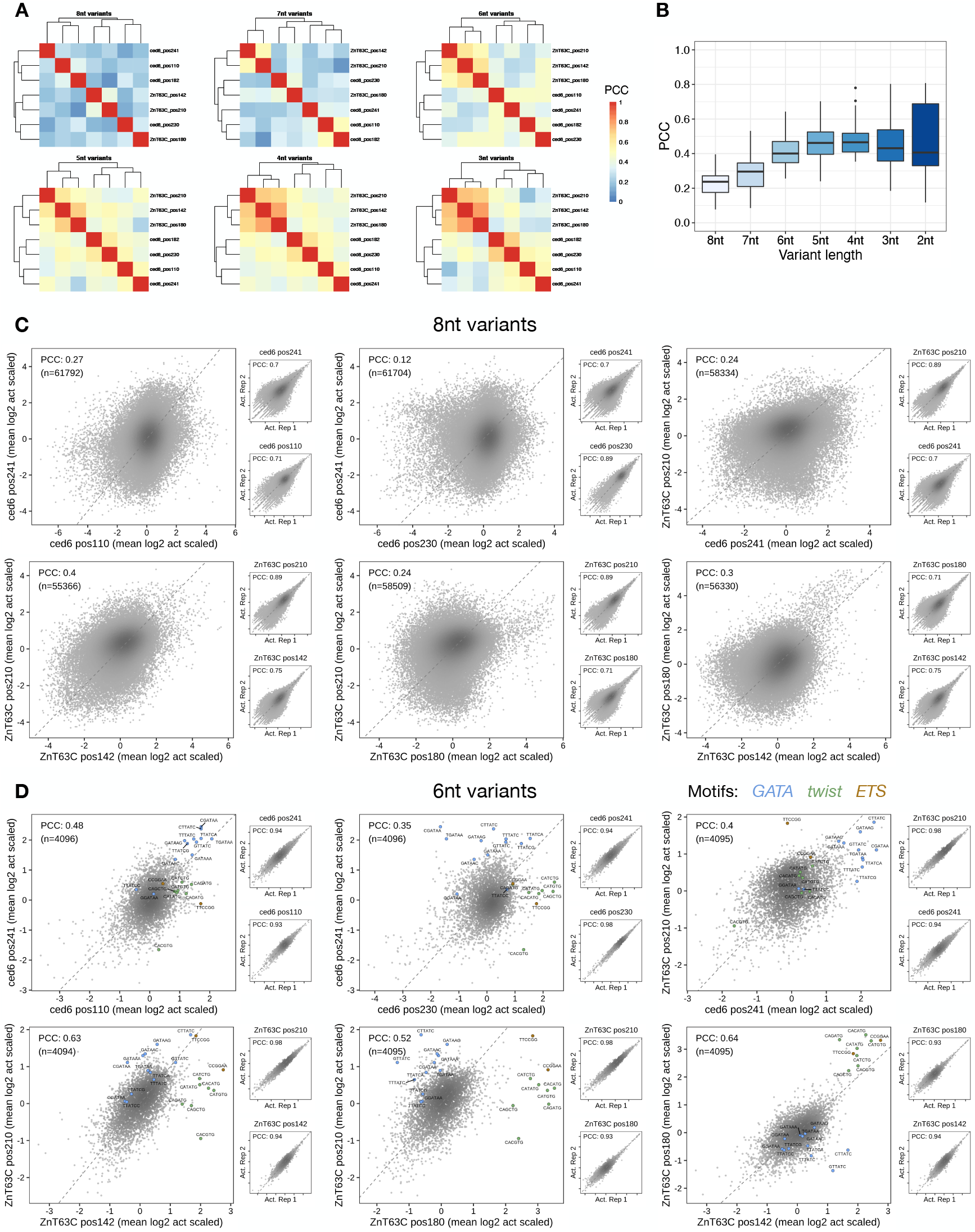
Comparison of all random variants across enhancer positions. **A)** Hierarchical clustering of all enhancer positions based on PCC of variant enhancer activities in each position, when considering different lengths of sequence variants (see Methods). **B)** Distribution of PCCs from (A) in function of the length of sequence variants considered. **C,D)** Comparison of z-scores of log2 enhancer activity of all 8nt **(C)** or 6nt **(D**; see Methods) variants between enhancer positions (insets show activity for replicates (Act. Rep) 1 versus 2 for each position). Color reflects the enhancer position and point density. PCCs and number of sequence variants are shown. Variants matching to GATA, twist and ETS motifs are highlighted in (D).

**Supplementary Figure 8.**
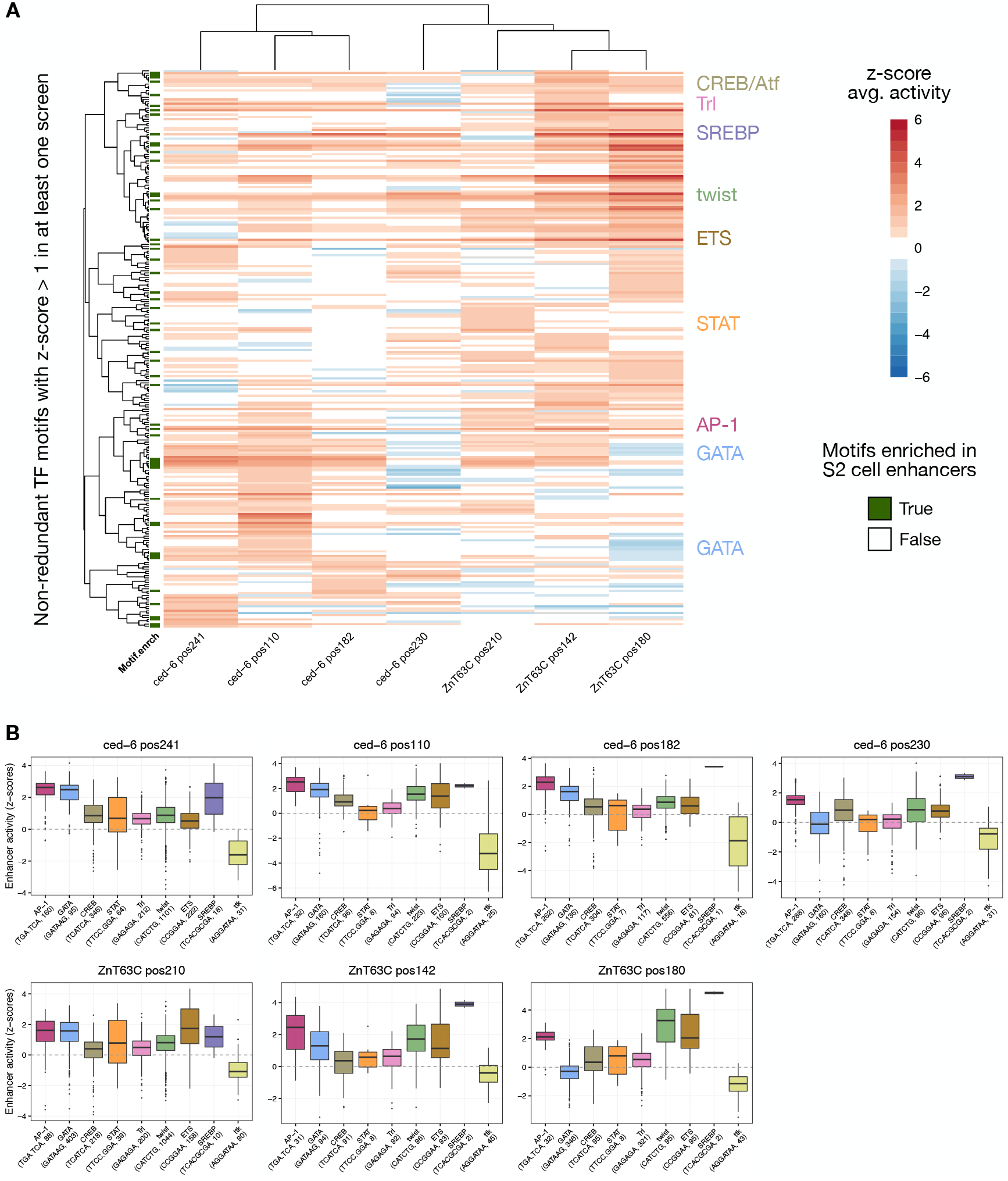
Activity of TF motif types at different enhancer positions. **A)** Heatmap of average z-scores of log2 enhancer activity of variants creating each TF motif type across all seven enhancer positions. Only motif types active (average z-score > 1) in at least one position are shown. Motifs and enhancer positions were clustered using hierarchical clustering and their activity is colored in shades of red (activating) and blue (repressing). Motifs enriched in S2 cell enhancers are labelled in green. Motif types used in the motif pasting experiment are highlighted. **B)** Activity of different TF motifs at each enhancer position. Distribution of z-scores of log2 enhancer activity for variants creating each TF motifs in *ced-6* and *ZnT63C* enhancer positions.

**Supplementary Figure 9.**
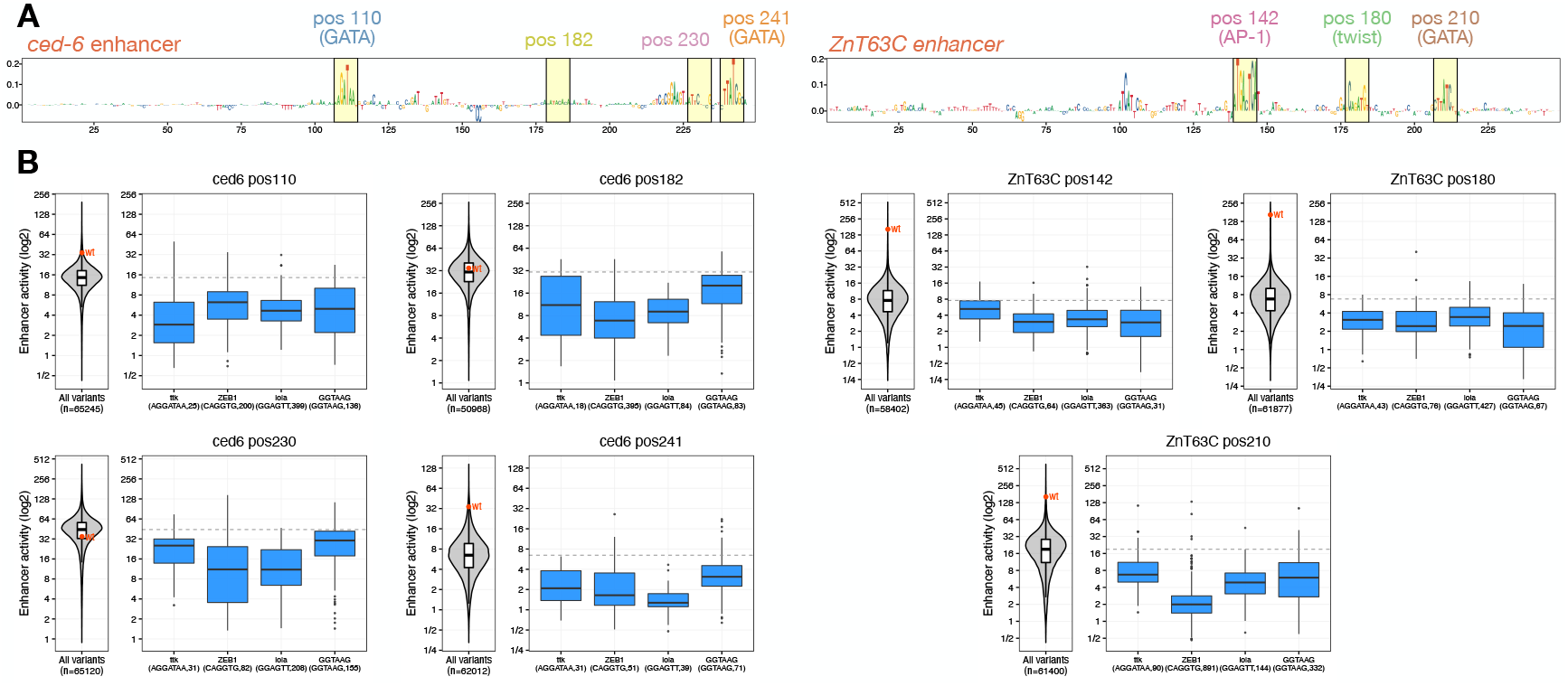
STARR-seq identifies known and novel motifs that repress enhancer activity. **A)** DeepSTARR-predicted nucleotide contribution scores for the *ced-6* (left) and *ZnT63C* (right) selected enhancer sequences. Selected 8nt motif positions and non-important control positions are highlighted in yellow with the respective numerical position, TF motif identity and different colors. **B)** Activity of different repressor motifs at each enhancer position. Distribution of enhancer activity for all enhancer variants (reft) or variants creating each repressor TF motif (right), per enhancer position. The activity of the wildtype sequence (wt, red) or median of all variants (grey dashed line) are shown. The string of each TF motif used for the motif matching and the number of variants matching to each motif are described in the x-axis: in the format “motif string (TF motif name, number of variants)”.

**Supplementary Figure 10.**
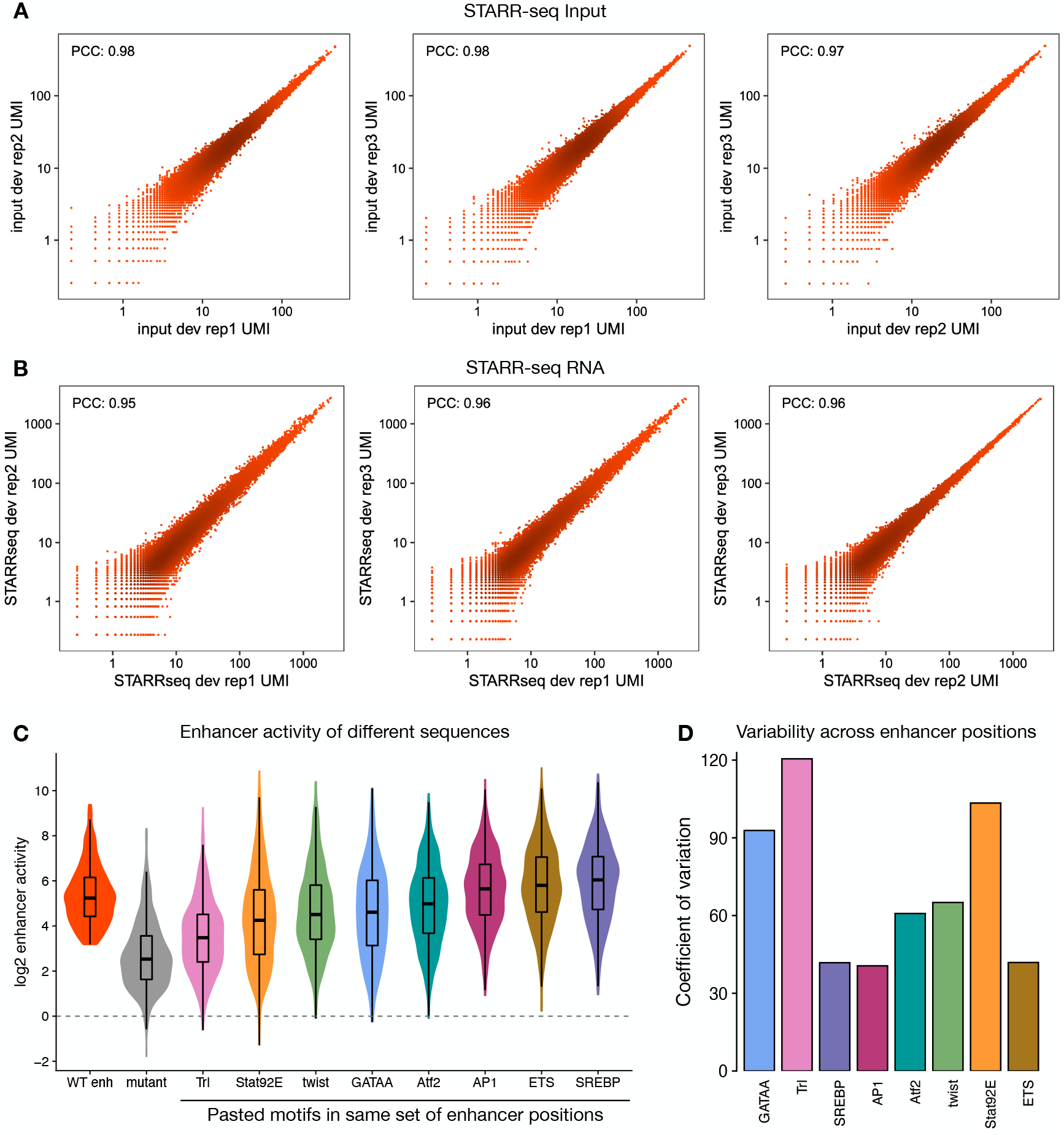
Systematic motif pasting screens in *Drosophila* enhancers. **A,B)** Pairwise comparisons of normalized STARR-seq input **(A)** and RNA **(B)** UMI read counts between three independent biological replicates across all oligos tested. Color reflects point density. The PCC is denoted for each comparison. **C)** Activity of pasted motifs at different enhancer positions. Distribution of enhancer activity changes (log2) of all wildtype enhancers used and their variants with either mutant sequences or different TF motifs pasted. **D)** Bar plots showing the coefficient of variation (ratio of the standard deviation to the mean) of the activity of each TF motif across all enhancer positions.

**Supplementary Figure 11.**
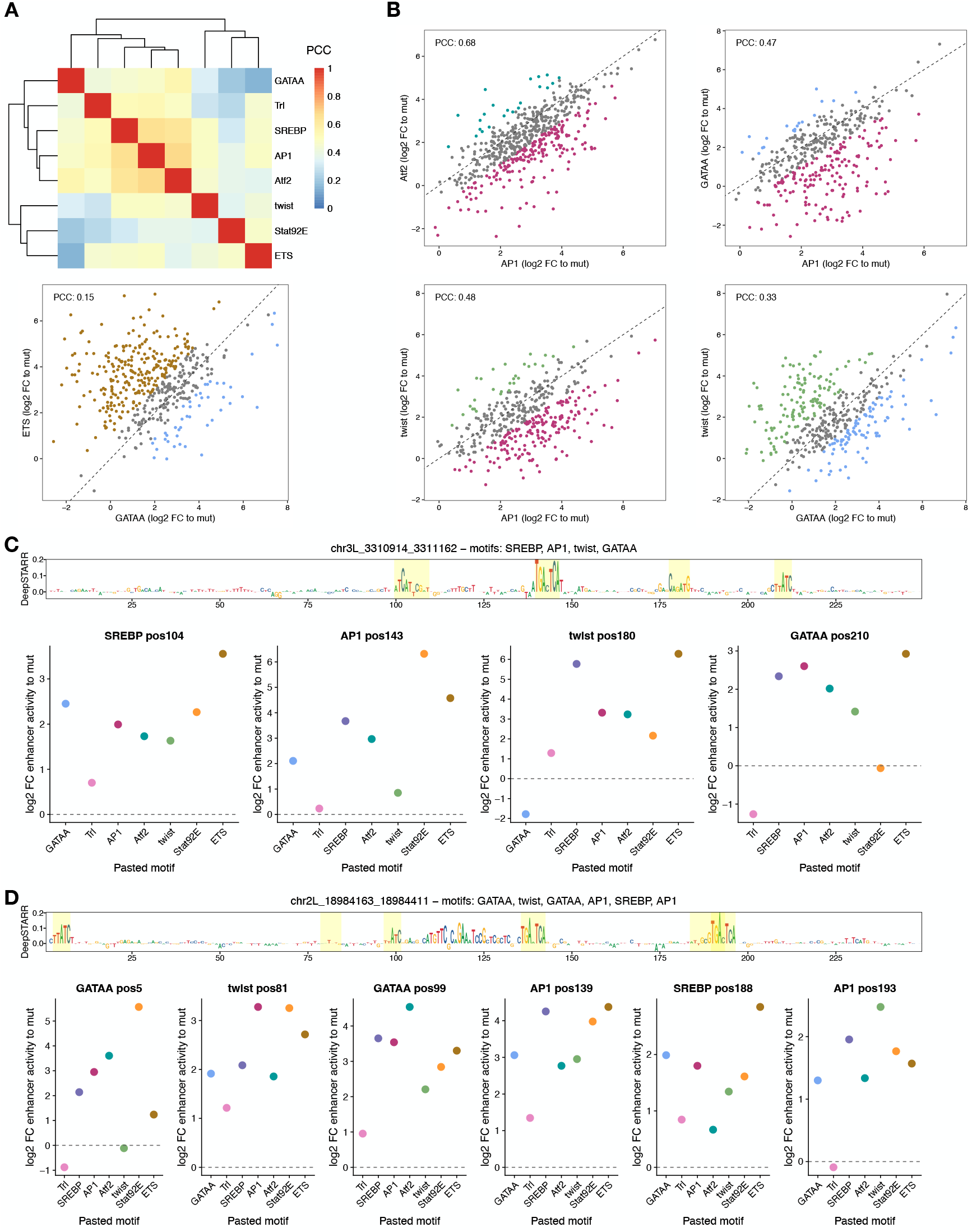
Motifs work differently at different enhancer positions. **A)** Hierarchical clustering of all TF motifs based on PCC of motif activities across all enhancer positions. **B)** Motifs work differently at different enhancer positions. Comparison between enhancer activity changes (log2 FC to mutated sequence) after pasting different TF motifs across all enhancer positions. Positions with stronger activity of each motif (>= 2-fold in respect to the other motif) are colored with the respective colors. PCC: Pearson correlation coefficient. **C,D)** DeepSTARR-predicted nucleotide contribution scores for two enhancers and respective positions (highlighted in yellow, with wildtype motif types described on top) included in the screen. For each position, the enhancer activity changes (log2 FC to mutated sequence) after pasting each TF motif are shown in dot plots (bottom).

**Supplementary Figure 12.**
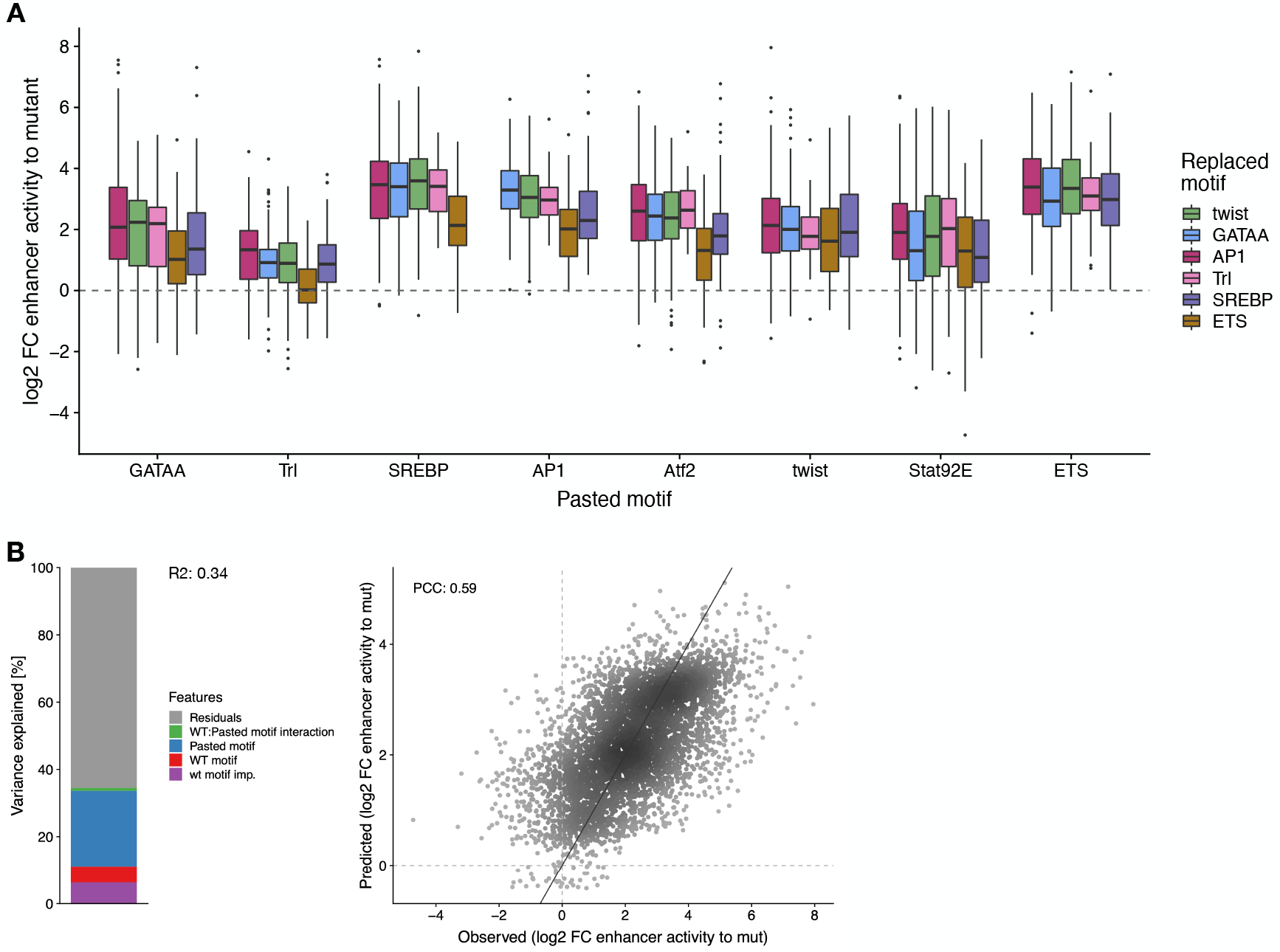
TF motif activity in function of wild type motif identity. **A)** Distribution of enhancer activity changes (log2 FC to mutated sequence) across all enhancer positions for each pasted TF motif, grouped by the identity of the wildtype motif. **B)** Left: Bar plot showing the amount of variance explained by the wildtype motif importance and identity, the pasted motif identity and the interaction between the wildtype and pasted motifs, using a linear model fit on all motif pasting results. Right: Scatter plots of predicted (linear model) vs. observed enhancer activity changes (log2 FC to mutated sequence) across all motif pasting experiments. Color reflects point density. PCC is shown.

**Supplementary Figure 13.**
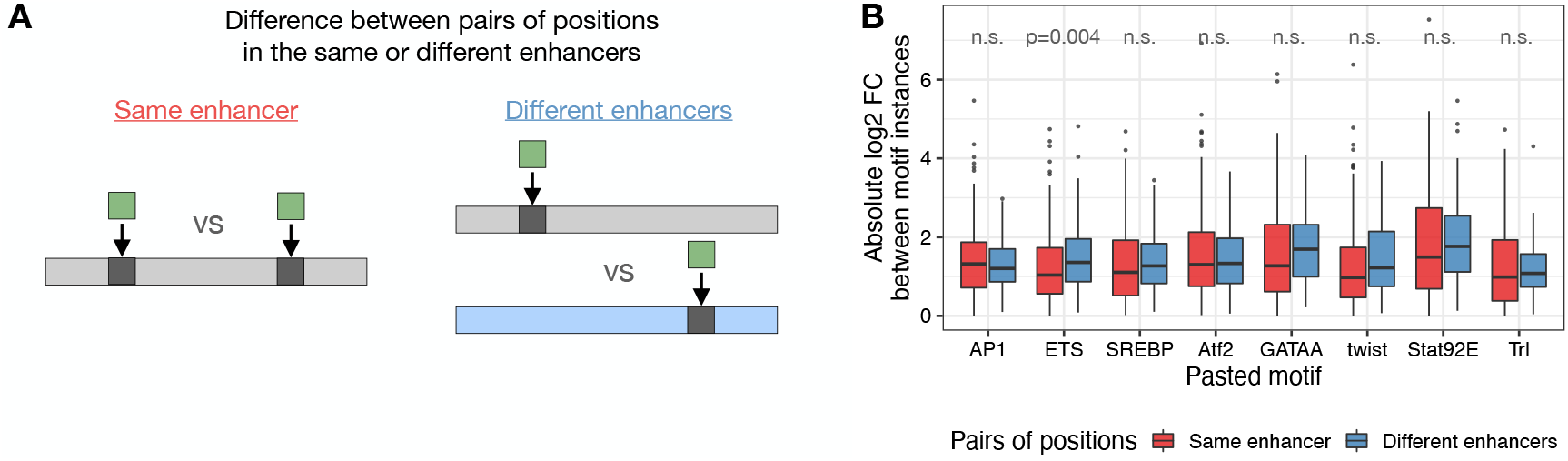
Motif activity in different positions in the same or different enhancers. **A)** Schematics of comparison of motif activity between instances within the same enhancer or in different enhancers. **B)** Absolute log2 fold-change in enhancer activity between instances within the same enhancer (red) or in different enhancers (blue) for each pasted TF motif type. n.s. non-significant (Wilcoxon signed rank test).

**Supplementary Figure 14.**
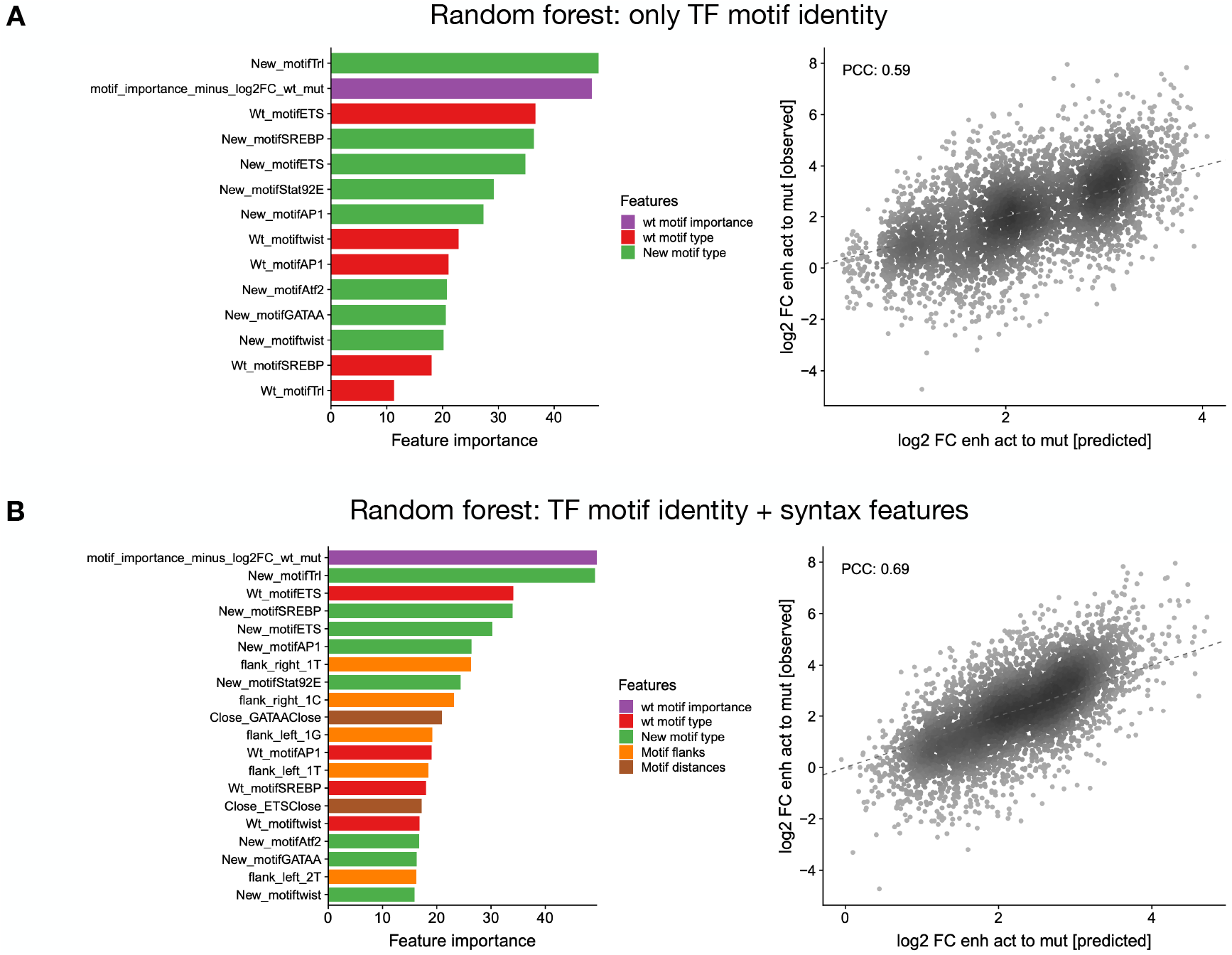
Prediction of motif activities using motif syntax features in random forest model. Left: Importance of all features **(A)** or only the top 20 **(B)** included in the random forest models with only TF motif identity **(A)** or also with syntax features **(B)**, sorted by importance and colored by feature type. Right: Scatter plots of predicted vs. observed enhancer activity changes (log2 FC to mutated sequence) across all motif pasting experiments. Color reflects point density. PCC is shown.

**Supplementary Figure 15.**
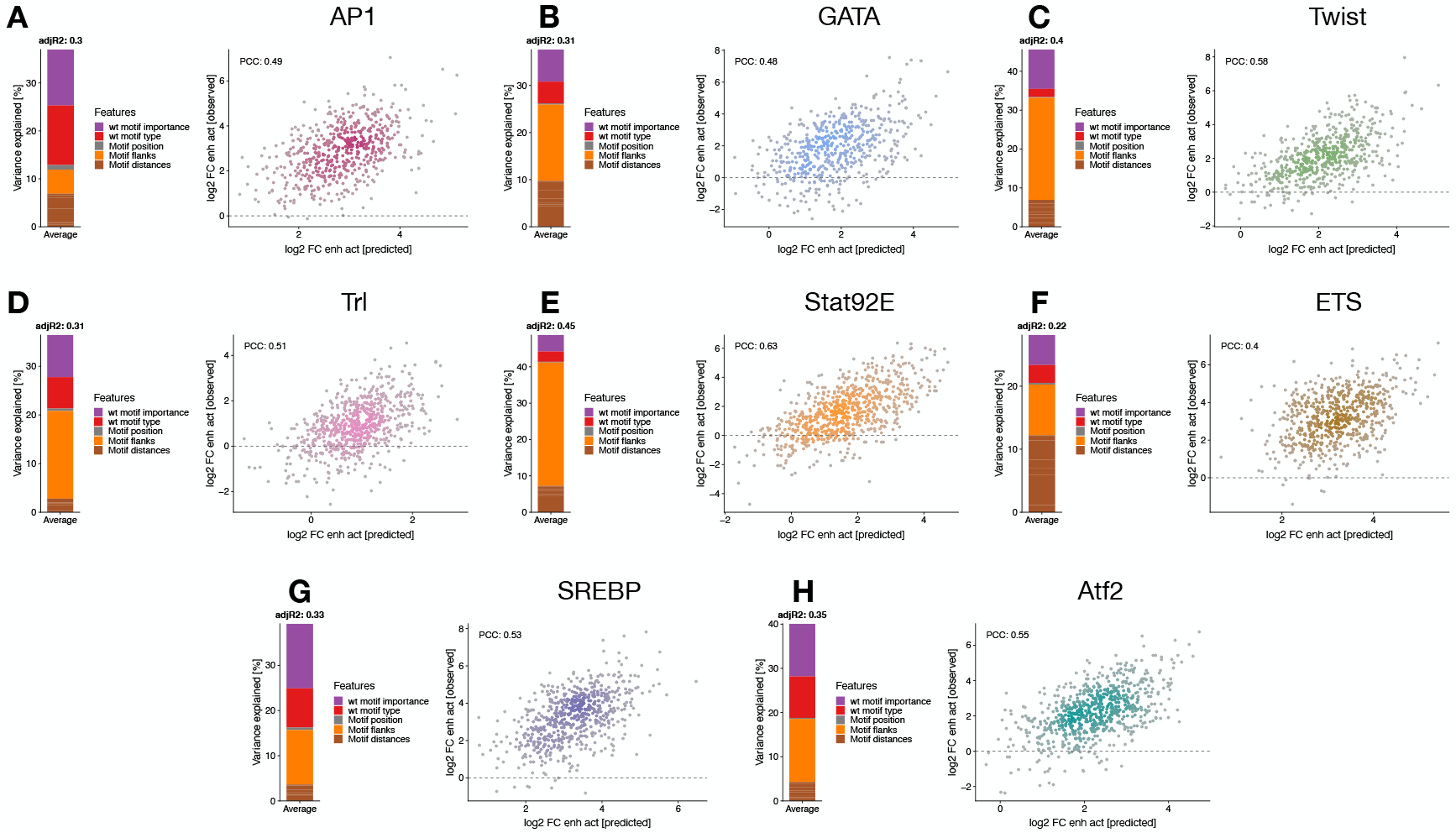
Linear models with syntax features to predict motif activities. **A-H)** Left: Bar plot showing the variance explained by the different types of features (color legend) for each of the linear models. Right: Scatter plots of predicted vs. observed enhancer activity changes (log2 FC to mutated sequence) for motif pasting experiments per TF motif type. Color reflects point density. PCC is shown.

**Supplementary Figure 16.**
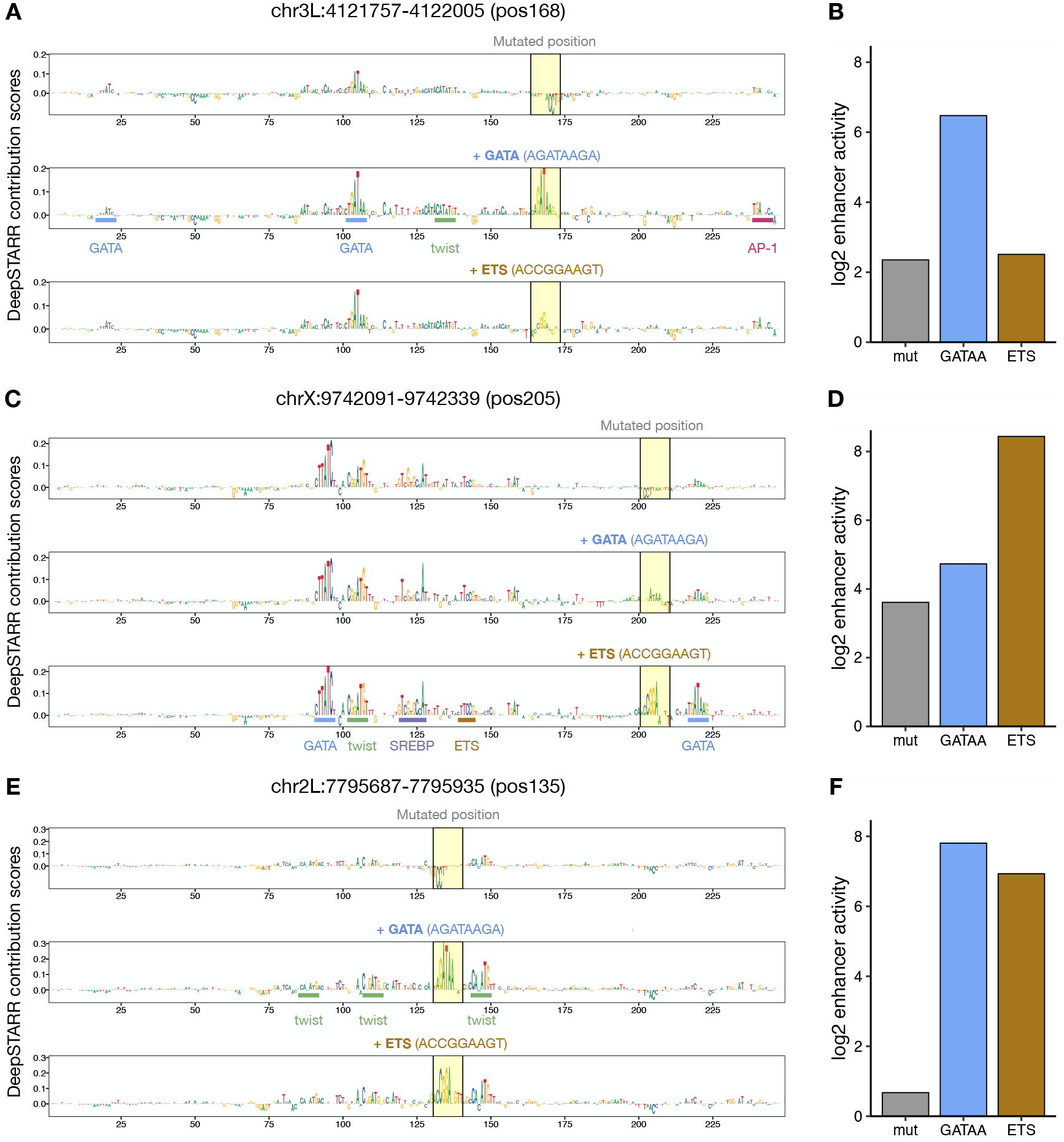
DeepSTARR-predicted importance scores for pasting GATA or ETS in the same positions. **A,C,E)** DeepSTARR-predicted nucleotide contribution scores for three different enhancers with a mutant sequence, GATA or ETS pasted at the highlighted positions. Motif sequences pasted are shown. **B,D,F)** Bar plots with enhancer activity (log2) of variants from (A,C,E).

**Supplementary Figure 17.**
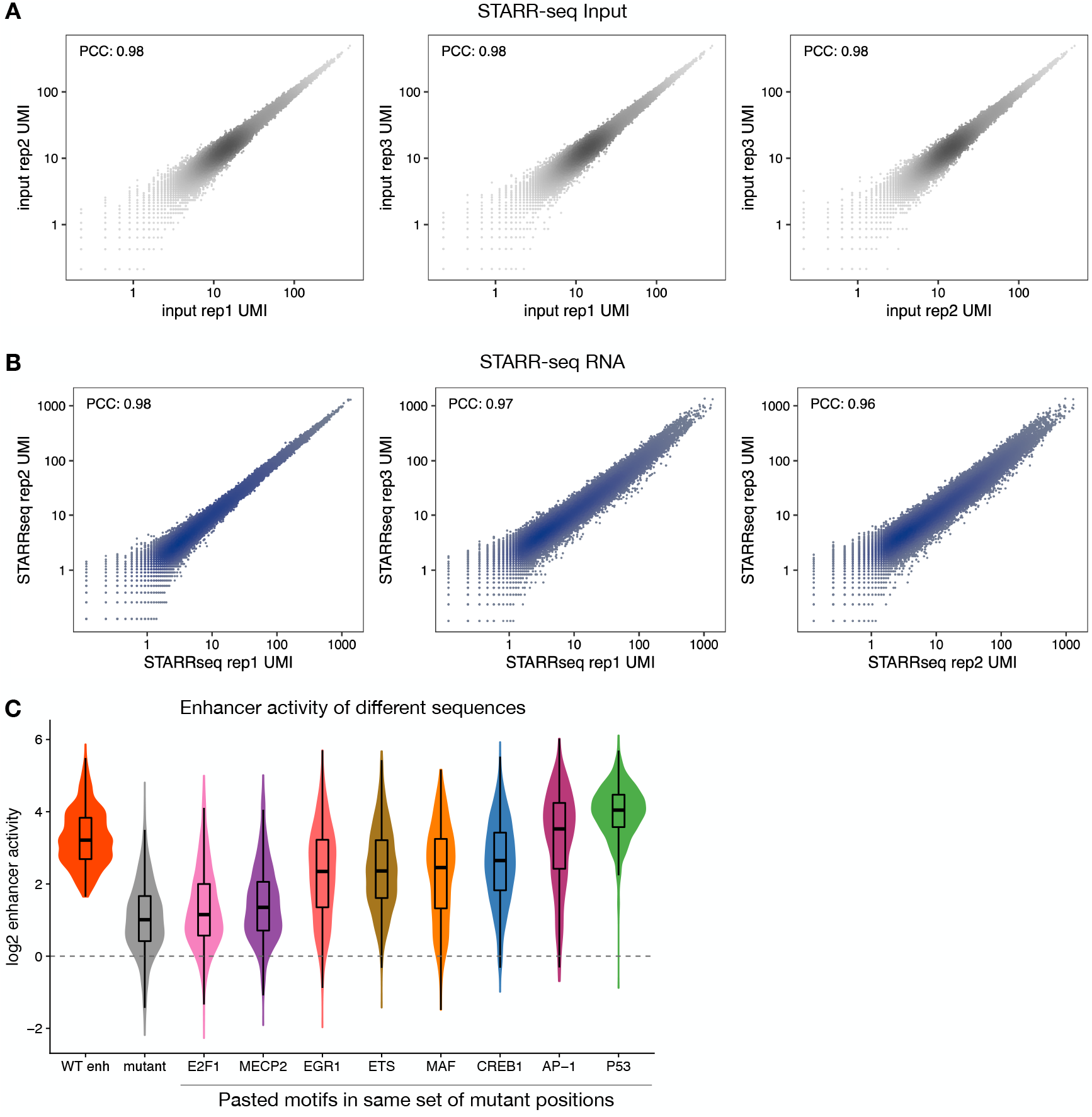
Systematic motif pasting screens in human enhancers. **A,B)** Pairwise comparisons of normalized STARR-seq input **(A)** and RNA **(B)** UMI read counts between three independent biological replicates across all oligos tested. Color reflects point density. The PCC is denoted for each comparison. **C)** Activity of pasted motifs at different enhancer positions. Distribution of enhancer activity changes (log2) of all wildtype enhancers used and their variants with either mutant sequences or different TF motifs pasted.

**Supplementary Figure 18.**
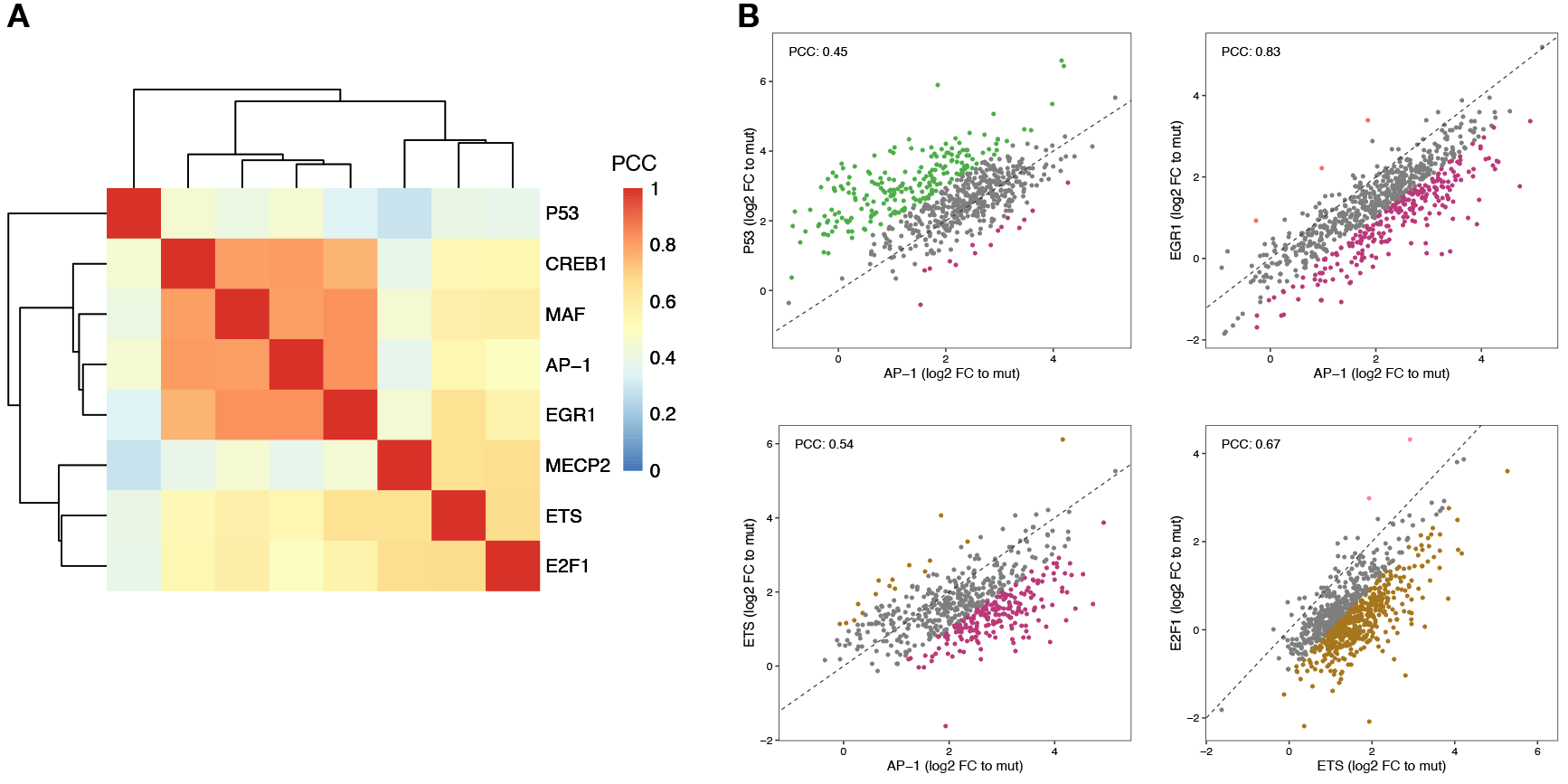
Human TF motifs work differently at different enhancer positions. **A)** Hierarchical clustering of all TF motifs based on PCC of motif activities across all enhancer positions. **B)** Motifs work differently at different enhancer positions. Comparison between enhancer activity changes (log2 FC to mutated sequence) after pasting different TF motifs across all enhancer positions. Positions with stronger activity of each motif (>= 2-fold in respect to the other motif) are colored with the respective colors. PCC: Pearson correlation coefficient.

**Supplementary Figure 19.**
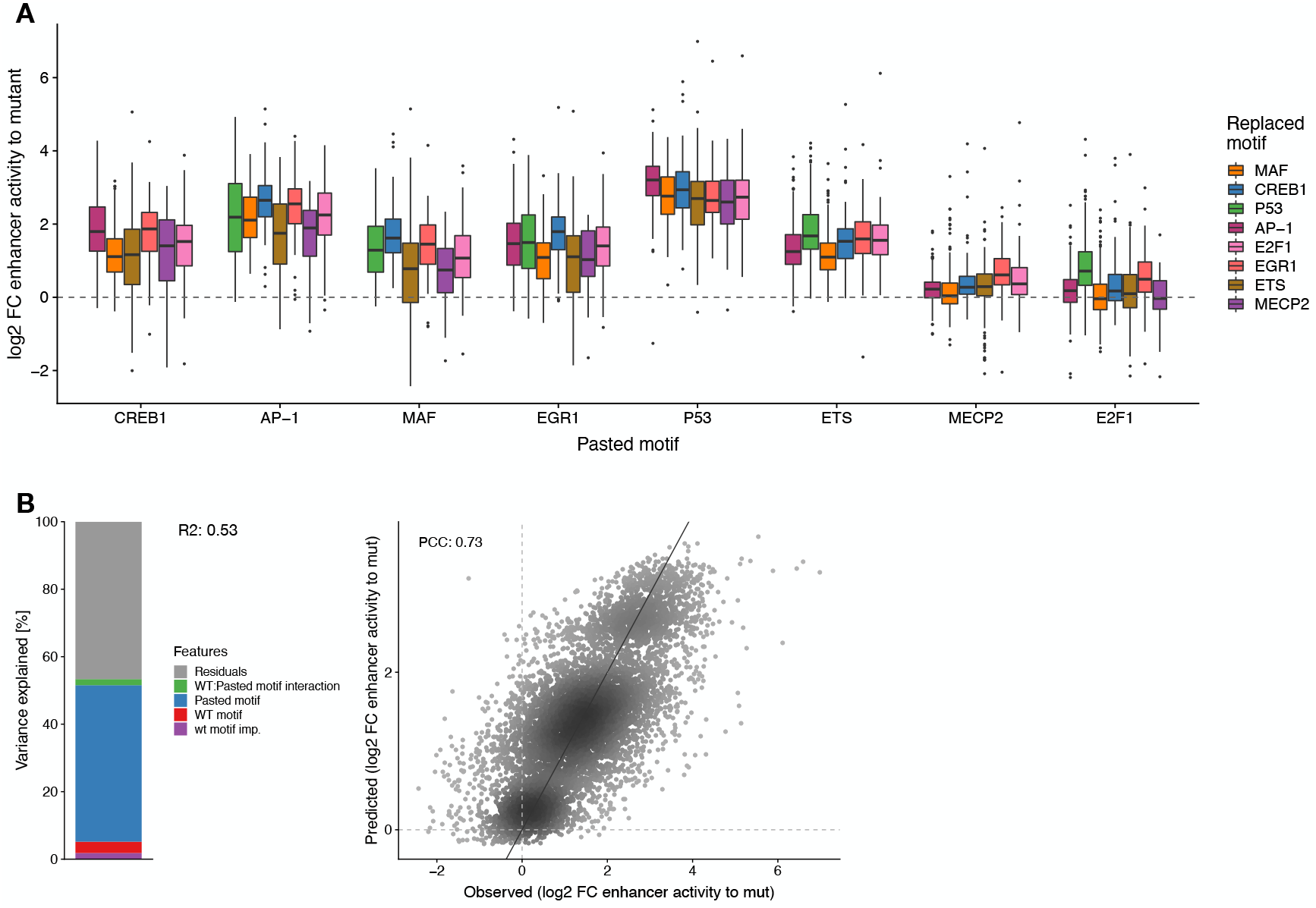
TF motif activity in function of wild type motif identity in human enhancers. **A)** Distribution of enhancer activity changes (log2 FC to mutated sequence) across all enhancer positions for each pasted TF motif, grouped by the identity of the wildtype motif. **B)** Left: Bar plot showing the amount of variance explained by the wildtype motif importance and identity, the pasted motif identity and the interaction between the wildtype and pasted motifs, using a linear model fit on all motif pasting results. Right: Scatter plots of predicted (linear model) vs. observed enhancer activity changes (log2 FC to mutated sequence) across all motif pasting experiments. Color reflects point density. PCC is shown.

**Supplementary Figure 20.**
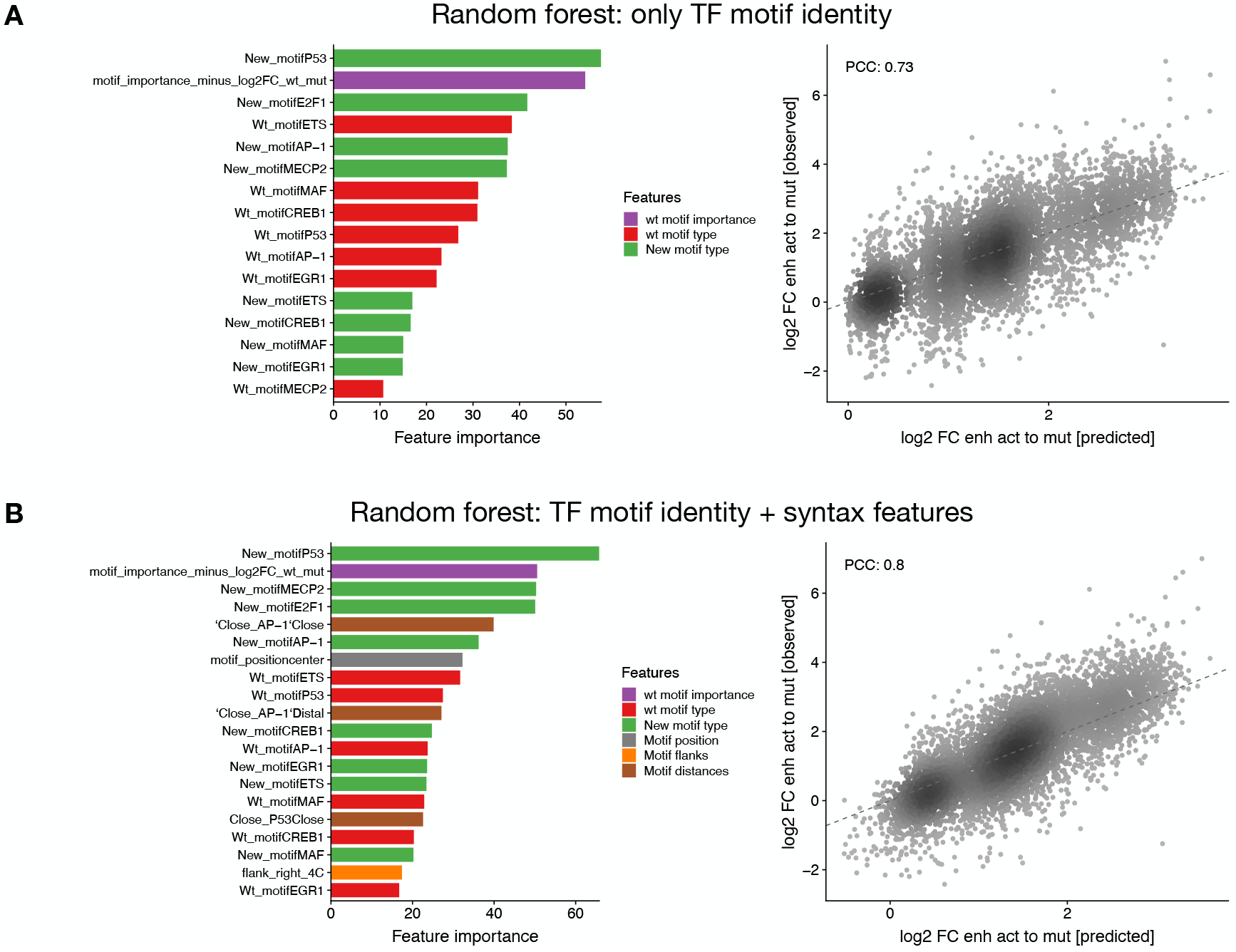
Prediction of motif activities using motif syntax features in human enhancers. Left: Importance of all features **(A)** or only the top 20 **(B)** included in the random forest models with only TF motif identity **(A)** or also with syntax features **(B)**, sorted by importance and colored by feature type. Right: Scatter plots of predicted vs. observed enhancer activity changes (log2 FC to mutated sequence) across all motif pasting experiments. Color reflects point density. PCC is shown.

**Supplementary Figure 21.**
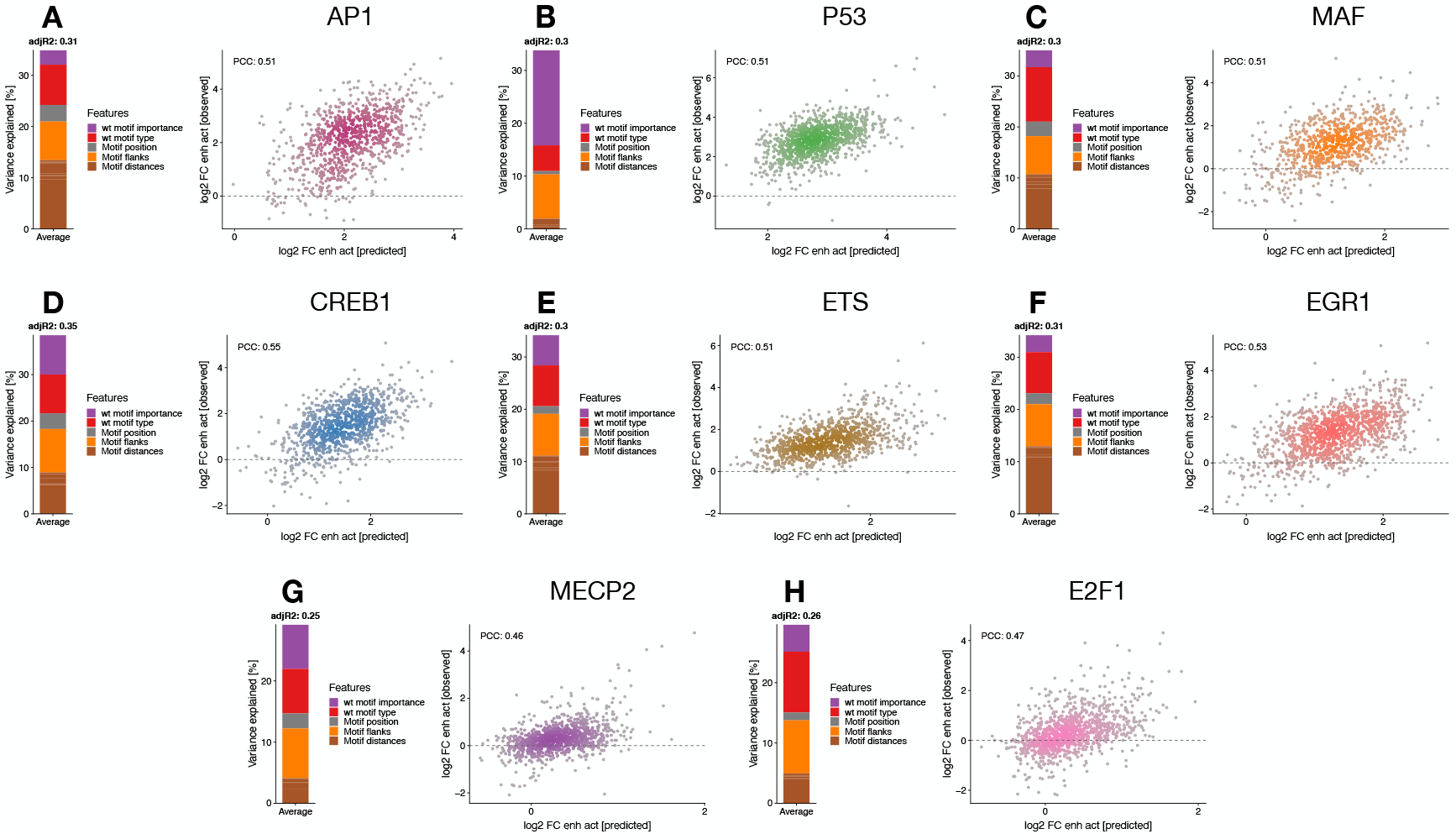
Linear models with syntax features to predict motif activities in human enhancers. **A-H)** Left: Bar plot showing the variance explained by the different types of features (color legend) for each of the linear models. Right: Scatter plots of predicted vs. observed enhancer activity changes (log2 FC to mutated sequence) for motif pasting experiments per TF motif type. Color reflects point density. PCC is shown.

**Supplementary Figure 22.**
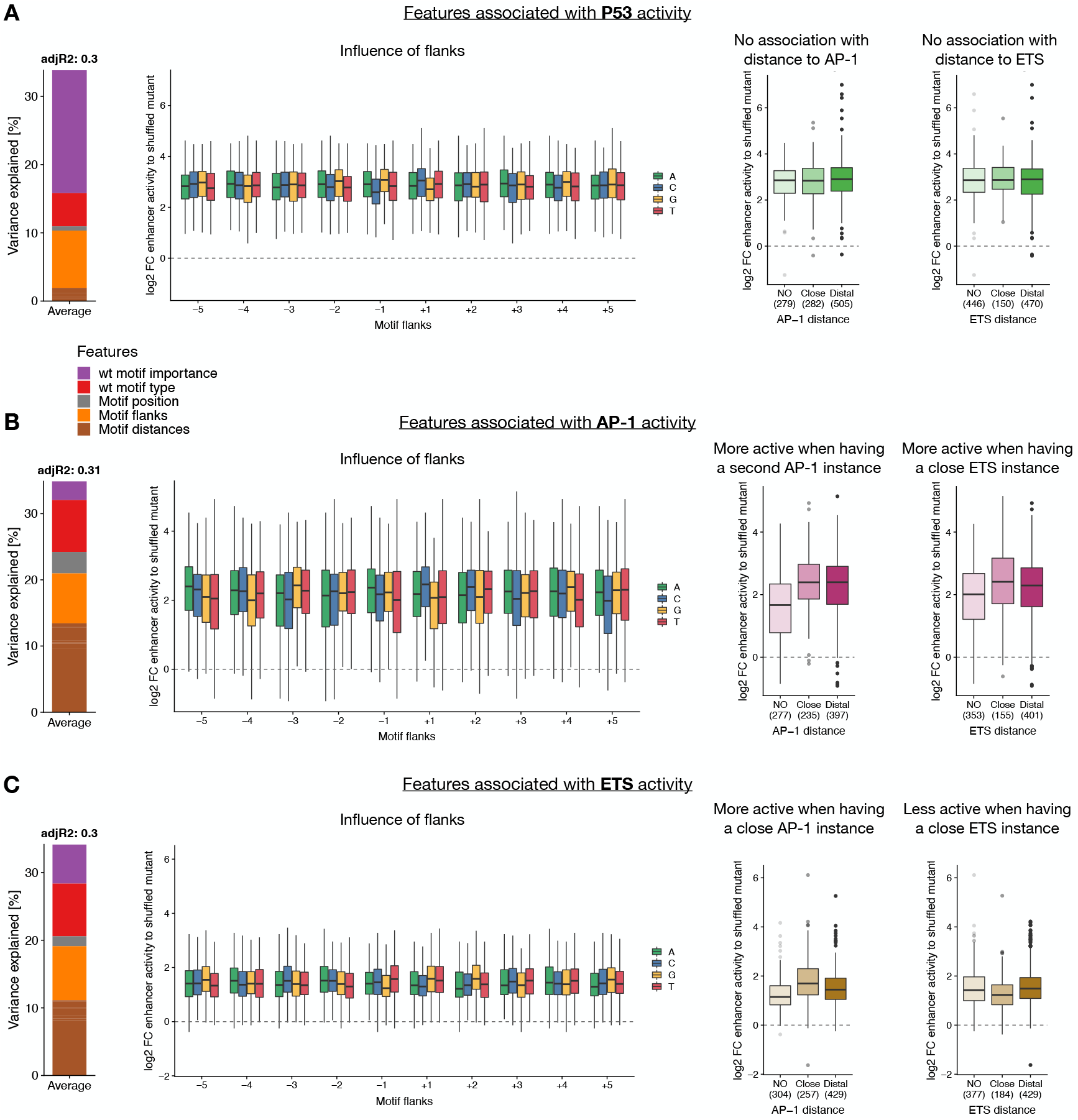
Sequence features associated with activity of P53, AP-1 and ETS motifs in human enhancers. **A-C)** Left: Bar plot showing the variance explained by the different types of features (color legend) for each of the linear models. Middle-left: Motif activity according to the different bases at each flanking position, colored by nucleotide identity. Middle-right and right: Enhancer activity changes (log2 FC to mutated sequence) after pasting each TF motif in positions with no additional AP-1 (middle-right) or ETS (right) in the enhancer, or with additional AP-1 or ETS at close (<= 25 bp) or distal (>25 bp) distances. Number of instances are shown.

**Supplementary Figure 23.**
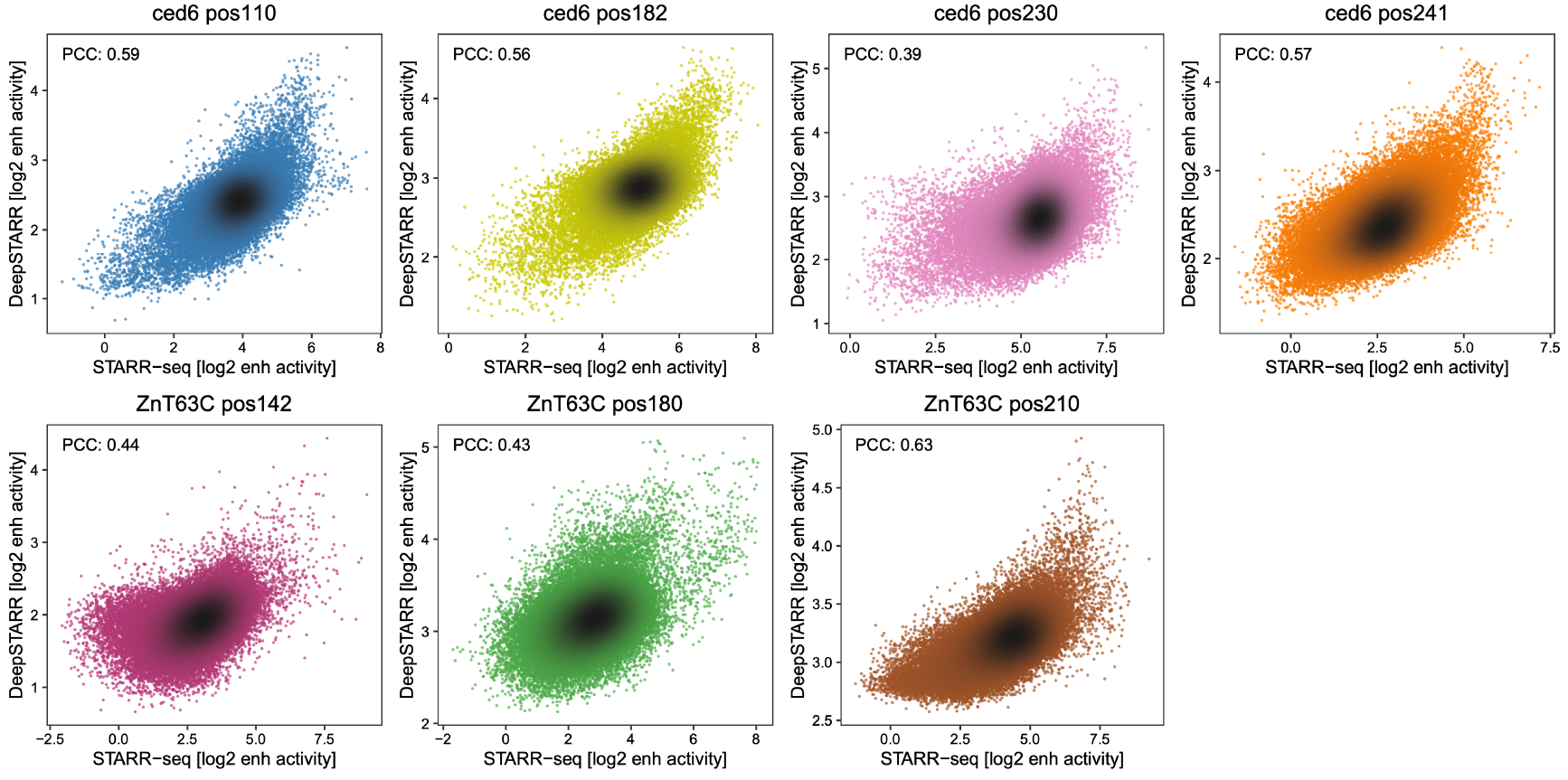
DeepSTARR predicts enhancer sequence changes. Comparison between DeepSTARR predicted (y-axis) and experimentally measured (x-axis) activity of random sequence variants tested at the different enhancer positions. Color reflects the enhancer position and point density. PCCs are shown.

**Supplementary Figure 24.**
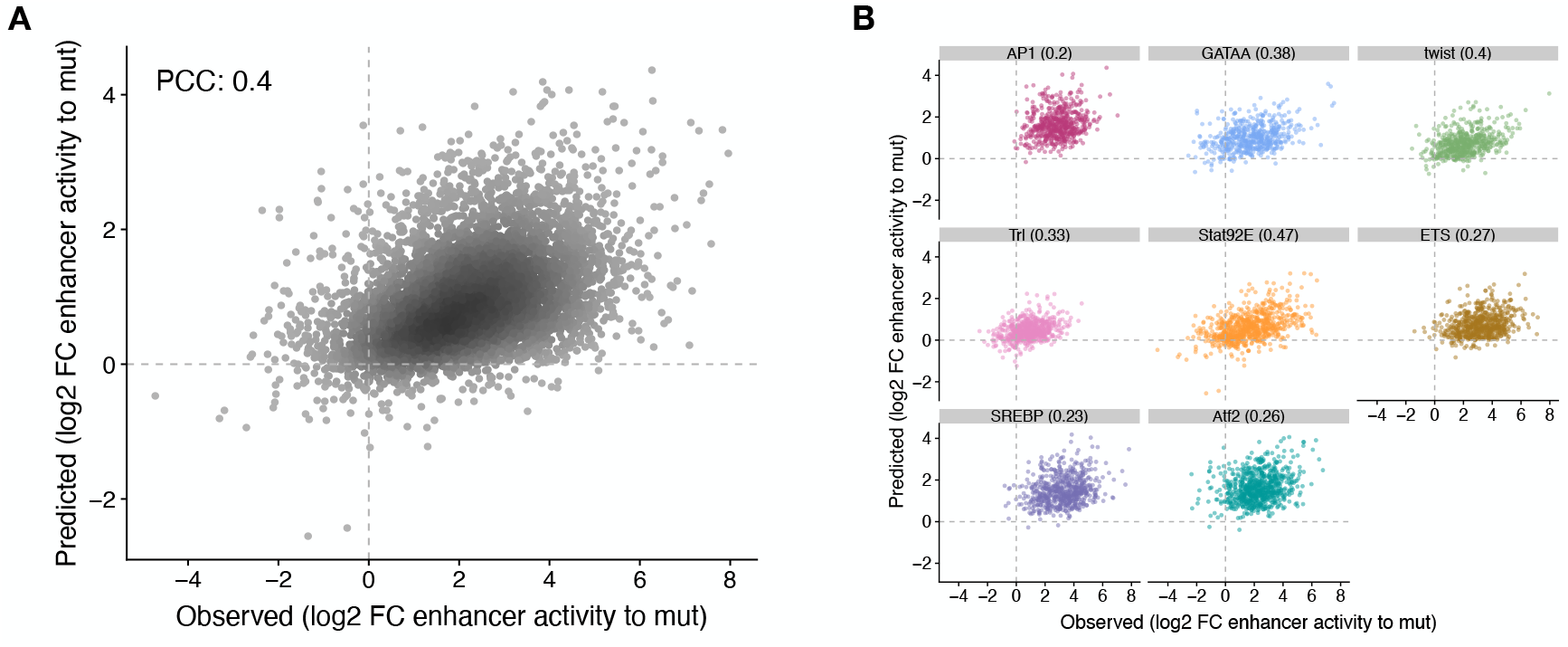
DeepSTARR predicts activity of motifs in different enhancer positions. **A)** Comparison between DeepSTARR predicted (y-axis) and experimentally measured (x-axis) enhancer activity changes (log2 FC to mutated sequence) for all motif pasting sequences. Color reflects the enhancer position and point density. PCCs are shown. **B)** Same as in (A) but per pasted TF motif.

## Supplementary Tables

**Supplementary Table 1. Primers used for UMI-STARR-seq library cloning.**

**Supplementary Table 2. Random variants and oligo UMI-STARR-seq mapping statistics.**

Summary of total sequenced reads, mapped reads and unique fragments (after collapsing by UMIs) for two random variants and three oligo UMI-STARR-seq screens in S2 cells, and three oligo UMI-STARR-seq screens in human HCT-116 cells.

**Supplementary Table 3. Activity of random variants in seven enhancer positions.**

8nt and 16nt forward and reverse sequences, activities and scaled activities in each of the seven enhancer positions.

**Supplementary Table 4. *Drosophila* and human TF motif sequences used in the motif pasting experiments.**

**Supplementary Table 5. Results of motif-pasting experiment in *Drosophila* S2 enhancers.**

Table with all oligos used in the analysis of *Drosophila* motif pasting with their DNA sequence, wildtype motif information, pasted motif information, activity of respective enhancer variant, of the original wildtype or motif-mutant enhancer, and respective log2 fold-changes.

**Supplementary Table 6. Results of motif-pasting experiment in human HCT-116 enhancers.**

Table with all oligos used in the analysis of human motif pasting with their DNA sequence, wildtype motif information, pasted motif information, activity of respective enhancer variant, of the original wildtype or motif-mutant enhancer, and respective log2 fold-changes.

## References

Arnold CD, Gerlach D, Spies D, Matts JA, Sytnikova YA, Pagani M, Lau NC, Stark A. 2014. Quantitative genome-wide enhancer activity maps for five Drosophila species show functional enhancer conservation and turnover during cis-regulatory evolution. Nat Genet 46: 685–692.

Arnold CD, Gerlach D, Stelzer C, Boryń ŁM, Rath M, Stark A. 2013. Genome-wide quantitative enhancer activity maps identified by STARR-seq. Science (1979) 339: 1074–1077.

Arnosti DN, Barolo S, Levine M, Small S. 1996. The eve stripe 2 enhancer employs multiple modes of transcriptional synergy. Development 122: 205–214.

Arnosti DN, Kulkarni MM. 2005. Transcriptional enhancers: Intelligent enhanceosomes or flexible billboards? J Cell Biochem 94: 890–898.

Avsec Ž, Weilert M, Shrikumar A, Krueger S, Alexandari A, Dalal K, Fropf R, Mcanany C, Gagneur J, Kundaje A, et al. 2021. Base-resolution models of transcription-factor binding reveal soft motif syntax. Nat Genet 53: 354–366.

Banerji J, Rusconi S, Schaffner W. 1981. Expression of a β-globin gene is enhanced by remote SV40 DNA sequences. Cell 27: 299–308.

Blow MJ, McCulley DJ, Li Z, Zhang T, Akiyama JA, Holt A, Plajzer-Frick I, Shoukry M, Wright C, Chen F, et al. 2010. ChIP-seq identification of weakly conserved heart enhancers. Nat Genet 42: 806–812.

Burda P, Laslo P, Stopka T. 2010. The role of PU.1 and GATA-1 transcription factors during normal and leukemogenic hematopoiesis. Leukemia 24: 1249–1257.

Crocker J, Abe N, Rinaldi L, McGregor AP, Frankel N, Wang S, Alsawadi A, Valenti P, Plaza S, Payre F, et al. 2015. Low Affinity Binding Site Clusters Confer Hox Specificityand Regulatory Robustness. Cell 160: 191–203.

de Almeida BP, Reiter F, Pagani M, Stark A. 2022. DeepSTARR predicts enhancer activity from DNA sequence and enables the de novo design of synthetic enhancers. Nat Genet.

de Boer CG, Vaishnav ED, Sadeh R, Abeyta EL, Friedman N, Regev A. 2019. Deciphering eukaryotic cis-regulatory logic with 100 million random promoters. Nat Biotechnol 38: 56–65.

Dror I, Golan T, Levy C, Rohs R, Mandel-Gutfreund Y. 2015. A widespread role of the motif environment in transcription factor binding across diverse protein families. Genome Res 25: 1268–1280.

Erceg J, Saunders TE, Girardot C, Devos DP, Hufnagel L, Furlong EEM. 2014. Subtle Changes in Motif Positioning Cause Tissue-Specific Effects on Robustness of an Enhancer’s Activity. PLoS Genet 10: e1004060.

Farley EK, Olson KM, Zhang W, Brandt AJ, Rokhsar DS, Levine MS. 2015. Suboptimization of developmental enhancers. Science (1979) 350: 325–328.

Farley EK, Olson KM, Zhang W, Rokhsar DS, Levine MS. 2016. Syntax compensates for poor binding sites to encode tissue specificity of developmental enhancers. Proceedings of the National Academy of Sciences 113: 6508–6513.

Fiore C, Cohen BA. 2016. Interactions between pluripotency factors specify cis-regulation in embryonic stem cells. Genome Res 26: 778–786.

Fisher S, Grice EA, Vinton RM, Bessling SL, McCallion AS. 2006. Conservation of RET regulatory function from human to zebrafish without sequence similarity. Science (1979) 312: 276–279.

Fuqua T, Jordan J, Breugel ME van, Halavatyi A, Tischer C, Polidoro P, Abe N, Tsai A, Mann RS, Stern DL, et al. 2020. Dense and pleiotropic regulatory information in a developmental enhancer. Nature 587: 235–239.

Galupa R, Alvarez-Canales G, Borst NO, Fuqua T, Gandara L, Misunou N, Richter K, Alves MRP, Karumbi E, Perkins ML, et al. 2022. Enhancer architecture and chromatin accessibility constrain phenotypic space during development. bioRxiv. https://doi.org/10.1101/2022.06.02.494376.

Gompel N, Prud’homme B, Wittkopp PJ, Kassner VA, Carroll SB. 2005. Chance caught on the wing: cis-regulatory evolution and the origin of pigment patterns in Drosophila. Nature 433: 481–487.

Hanes SD, Riddihough G, Ish-Horowicz D, Brent R. 1994. Specific DNA Recognition and Intersite Spacing Are Critical for Action of the Bicoid Morphogent. Mol Cell Biol 14: 3364–3375.

He Q, Bardet AF, Patton B, Purvis J, Johnston J, Paulson A, Gogol M, Stark A, Zeitlinger J. 2011. High conservation of transcription factor binding and evidence for combinatorial regulation across six Drosophila species. Nat Genet 43: 414–421.

Heinz S, Benner C, Spann N, Bertolino E, Lin YC, Laslo P, Cheng JX, Murre C, Singh H, Glass CK. 2010. Simple Combinations of Lineage-Determining Transcription Factors Prime cis-Regulatory Elements Required for Macrophage and B Cell Identities. Mol Cell 38: 576–589.

Janssens J, Aibar S, Taskiran II, Ismail JN, Spanier KI, Gonzalez-Blas CB, Quan XJ, Papasokrati D, Hulselmans G, Makhzami S, et al. 2022. Decoding gene regulation in the fly brain. Nature 601: 630–636.

Jindal GA, Farley EK. 2021. Enhancer grammar in development, evolution, and disease: dependencies and interplay. Dev Cell 56: 575–587.

King DM, Hong CKY, Shepherdson JL, Granas DM, Maricque BB, Cohen BA. 2020. Synthetic and genomic regulatory elements reveal aspects of cis-regulatory grammar in mouse embryonic stem cells. Elife 9: 1–24.

Kolde R. 2019. pheatmap: Pretty Heatmaps. R package version 1.0.12. https://CRAN.R-project.org/package=pheatmap.

Kuhn M. 2018. caret: Classification and Regression Training. R package version 6.0–80. https://CRAN.R-project.org/package=caret.

Kulkarni MM, Arnosti DN. 2003. Information display by transcriptional enhancers. Development 130: 6569–6575.

Kvon EZ, Kazmar T, Stampfel G, Yáñez-Cuna JO, Pagani M, Schernhuber K, Dickson BJ, Stark A. 2014. Genome-scale functional characterization of Drosophila developmental enhancers in vivo. Nature 512: 91–95.

Langmead B, Trapnell C, Pop M, Salzberg SL. 2009. Ultrafast and memory-efficient alignment of short DNA sequences to the human genome. Genome Biol 10: R25.

Levine M. 2010. Transcriptional Enhancers in Animal Development and Evolution. Current Biology 20: R754–R763.

Li R, Pei H, Watson DK. 2000. Regulation of Ets function by protein – Protein interactions. Oncogene 19: 6514–6523.

Liu F, Posakony JW. 2012. Role of architecture in the function and specificity of two notch-regulated transcriptional enhancer modules. PLoS Genet 8: e1002796.

Love MI, Huber W, Anders S. 2014. Moderated estimation of fold change and dispersion for RNA-seq data with DESeq2. Genome Biol 15: 1–21.

Ludwig MZ, Bergman C, Patel NH, Kreitman M. 2000. Evidence for stabilizing selection in a eukaryotic enhancer element. Nature 403: 564–567.

Ludwig MZ, Patel NH, Kreitman M. 1998. Functional analysis of eve stripe 2 enhancer evolution in Drosophila: rules governing conservation and change. Development 125.

Lundberg SM, Erion G, Chen H, DeGrave A, Prutkin JM, Nair B, Katz R, Himmelfarb J, Bansal N, Lee S-I. 2020. From local explanations to global understanding with explainable AI for trees. Nat Mach Intell 2: 56–67.

Lundberg SM, Lee S-I. 2017. A Unified Approach to Interpreting Model Predictions. 31st Conference on Neural Information Processing Systems.

Mathelier A, Xin B, Chiu TP, Yang L, Rohs R, Wasserman WW. 2016. DNA Shape Features Improve Transcription Factor Binding Site Predictions In Vivo. Cell Syst 3: 278–286.

May D, Blow MJ, Kaplan T, McCulley DJ, Jensen BC, Akiyama JA, Holt A, Plajzer-Frick I, Shoukry M, Wright C, et al. 2012. Large-scale discovery of enhancers from human heart tissue. Nat Genet 44: 89–93.

Muerdter F, Boryn ŁM, Woodfin AR, Neumayr C, Rath M, Zabidi MA, Pagani M, Haberle V, Kazmar T, Catarino RR, et al. 2018. Resolving systematic errors in widely used enhancer activity assays in human cells. Nat Methods 15: 141–149.

Neumayr C, Pagani M, Stark A, Arnold CD. 2019. STARR-seq and UMI-STARR-seq: Assessing Enhancer Activities for Genome-Wide-, High-, and Low-Complexity Candidate Libraries. Curr Protoc Mol Biol 128: e105.

Omar Wagih. 2017. ggseqlogo: A “ggplot2” Extension for Drawing Publication-Ready Sequence Logos. R package version 0.1. https://CRAN.R-project.org/package=ggseqlogo.

Panne D. 2008. The enhanceosome. Curr Opin Struct Biol 18: 236–242.

R Core Team. 2020. R: A language and environment for statistical computing. R Foundation for Statistical Computing, Vienna, Austria. URL https://www.R-project.org/.

Rastegar S, Hess I, Dickmeis T, Nicod JC, Ertzer R, Hadzhiev Y, Thies WG, Scherer G, Strähle U. 2008. The words of the regulatory code are arranged in a variable manner in highly conserved enhancers. Dev Biol 318: 366–377.

Reiter F, Wienerroither S, Stark A. 2017. Combinatorial function of transcription factors and cofactors. Curr Opin Genet Dev 43: 73–81.

Rickels R, Shilatifard A. 2018. Enhancer Logic and Mechanics in Development and Disease. Trends Cell Biol 28: 608–630.

Sarkisyan KS, Bolotin DA, Meer M v., Usmanova DR, Mishin AS, Sharonov G v., Ivankov DN, Bozhanova NG, Baranov MS, Soylemez O, et al. 2016. Local fitness landscape of the green fluorescent protein. Nature 533: 397–401.

Schep A. 2021. motifmatchr: Fast Motif Matching in R. R package version 114.0.

Schmidt D, Wilson MD, Ballester B, Schwalie PC, Brown GD, Marshall A, Kutter C, Watt S, Martinez-Jimenez CP, Mackay S, et al. 2010. Five-Vertebrate ChIP-seq Reveals the Evolutionary Dynamics of Transcription Factor Binding. Science (1979) 1036–1040.

Schneider TD, Stephens RM. 1990. Sequence logos: a new way to display consensus sequences. Nucleic Acids Res 18: 6097.

Schneider TD, Stormo GD, Gold L, Ehrenfeucht A. 1986. Information content of binding sites on nucleotide sequences. J Mol Biol 188: 415–431.

Sharon E, Kalma Y, Sharp A, Raveh-Sadka T, Levo M, Zeevi D, Keren L, Yakhini Z, Weinberger A, Segal E. 2012. Inferring gene regulatory logic from high-throughput measurements of thousands of systematically designed promoters. Nat Biotechnol 30: 521–530.

Shrikumar A, Greenside P, Kundaje A. 2017. Learning important features through propagating activation differences. ArXiv 1704.02685.

Smith RP, Taher L, Patwardhan RP, Kim MJ, Inoue F, Shendure J, Ovcharenko I, Ahituv N. 2013. Massively parallel decoding of mammalian regulatory sequences supports a flexible organizational model. Nat Genet 45: 1021–1028.

Somermeyer LG, Fleiss A, Mishin AS, Bozhanova NG, Igolkina AA, Meiler J, Alaball Pujol M-E, Putintseva E v, Sarkisyan KS, Kondrashov FA. 2022. Heterogeneity of the GFP fitness landscape and data-driven protein design. Elife 11.

Spitz F, Furlong EEM. 2012. Transcription factors: From enhancer binding to developmental control. Nat Rev Genet 13: 613–626.

Swanson CI, Evans NC, Barolo S. 2010. Structural Rules and Complex Regulatory Circuitry Constrain Expression of a Notch- and EGFR-Regulated Eye Enhancer. Dev Cell 18: 359–376.

Swanson CI, Schwimmer DB, Barolo S. 2011. Rapid evolutionary rewiring of a structurally constrained eye enhancer. Current Biology 21: 1186–1196.

Taher L, McGaughey DM, Maragh S, Aneas I, Bessling SL, Miller W, Nobrega MA, McCallion AS, Ovcharenko I. 2011. Genome-wide identification of conserved regulatory function in diverged sequences. Genome Res 21: 1139–1149.

Thanos D, Maniatis T. 1995. Virus induction of human IFNβ gene expression requires the assembly of an enhanceosome. Cell 83: 1091–1100.

Vaishnav ED, de Boer CG, Molinet J, Yassour M, Fan L, Adiconis X, Thompson DA, Levin JZ, Cubillos FA, Regev A. 2022. The evolution, evolvability and engineering of gene regulatory DNA. Nature 603: 455–463.

Verfaillie A, Svetlichnyy D, Imrichova H, Davie K, Fiers M, Atak ZK, Hulselmans G, Christiaens V, Aerts S. 2016. Multiplex enhancer-reporter assays uncover unsophisticated TP53 enhancer logic. Genome Res 26: 882–895.

Villar D, Berthelot C, Aldridge S, Rayner TF, Lukk M, Pignatelli M, Park TJ, Deaville R, Erichsen JT, Jasinska AJ, et al. 2015. Enhancer evolution across 20 mammalian species. Cell 160: 554–566.

Visel A, Rubin EM, Pennacchio LA. 2009. Genomic views of distant-acting enhancers. Nature 461: 199–205.

Vockley CM, McDowell IC, D’Ippolito AM, Reddy TE. 2017. A long-range flexible billboard model of gene activation. Transcription 8: 261–267.

Weirauch MT, Hughes TR. 2010. Conserved expression without conserved regulatory sequence: the more things change, the more they stay the same. Trends in Genetics 26: 66–74.

Wickham H. 2016. ggplot2: Elegant Graphics for Data Analysis. Springer-Verlag New York. ISBN 978-3-319-24277-4, http://ggplot2.org.

Wong ES, Zheng D, Tan SZ, Bower NI, Garside V, Vanwalleghem G, Gaiti F, Scott E, Hogan BM, Kikuchi K, et al. 2020. Deep conservation of the enhancer regulatory code in animals. Science (1979) 370.

Xiong W-C, Montell C. 1993. tramtrack is a transcriptional repressor required for cell fate determination in the Drosophila eye. Genes Dev 7: 1085–1096.

Zabidi MA, Arnold CD, Schernhuber K, Pagani M, Rath M, Frank O, Stark A. 2015. Enhancer-core-promoter specificity separates developmental and housekeeping gene regulation. Nature 518: 556–559.

Zinzen RP, Senger K, Levine M, Papatsenko D. 2006. Computational Models for Neurogenic Gene Expression in the Drosophila Embryo. Current Biology 16: 1358– 1365.

